# Cell-specific imputation of drug connectivity mapping with incomplete data

**DOI:** 10.1101/2020.08.10.231720

**Authors:** Diana Sapashnik, Rebecca Newman, Christopher Michael Pietras, Fangfang Qu, Lior Kofman, Sean Boudreau, Inbar Fried, Donna K. Slonim

## Abstract

**Motivation:** Drug repositioning allows expedited discovery of new applications for existing compounds, but re-screening vast compound libraries is often prohibitively expensive. “Connectivity mapping” is a process that links drugs to diseases by identifying compounds whose impact on expression in a collection of cells reverses the disease’s impact on expression in disease-relevant tissues. The high throughput LINCS project has expanded the universe of compounds and cell types for which data are available, but even with this effort, many potentially clinically useful combinations are missing. To evaluate the possibility of repurposing drugs this way despite missing data, we compared collaborative filtering with either neighborhood-based or SVD imputation methods to two naive approaches via cross-validation.

**Results:** Methods were evaluated for their ability to predict drug connectivity despite missing data. Predictions improved when cell type was taken into account. Neighborhood-based collaborative filtering was the most successful method, with the best improvements in non-immortalized primary cells. We also explored which classes of compounds are most and least reliant on cell type for accurate imputation, and we identified connections between related compounds even when many were not measured in the relevant cells. We conclude that even for cells in which drug responses have not been fully characterized, it is possible to identify unassayed drugs that reverse in those cells the expression signatures observed in disease.

## 1 INTRODUCTION

*Connectivity mapping* (Lamb *et al.*, 2006) refers to the process of drug repositioning by finding candidate drugs that best reverse the expression changes caused by a given disease or condition. The original Connectivity Map database used microarrays to profile gene expression changes in up to four cancer cell lines treated with 164 perturbagens, many of them FDA-approved drugs. An expression “signature” from a given disease state, essentially sets of up- and down-regulated genes in relevant tissues from patients with the disease compared to normal controls, could then be used to find database compounds whose effect on gene expression was negatively correlated with the expression changes caused by the disease. Some compounds identified in this way were shown in further studies to have high potential for therapeutic efficacy in disease (Lamb *et al.*, 2006; Wei *et al.*, 2006; Zhang *et al.*, 2012).

More recently, the LINCS Consortium dramatically scaled up the connectivity map database using the L1000 assay, which measures expression of 978 genes at a much lower cost. The LINCS connectivity map includes a much larger set of compounds, small molecules, and cellular perturbations across a wider range of cell types (Subramanian *et al.*, 2017). However, while there are now many more cells profiled in the connectivity database, the data matrix is still very sparse, with most drug profiles in a small set of cancer cell lines. Yet recent work has shown that even different breast cancer cell lines can have different, context-specific responses to perturbation (Niepel *et al.*, 2017). We observe that variation in primary cells’ responses is even greater.

Further, gene expression profiles in primary cells are a relatively small part of the LINCS data set, and profiling all possible cell types and states is impractical even with more efficient assays. Thus, candidate drugs identified through connectivity mapping may have very different effects *in vivo*. For a precision medicine approach to connectivity, particularly outside the realm of oncology, the ability to identify drugs that reverse a patient’s disease signature even in the absence of connectivity data for a given cell/drug combination will improve scalability and relevance. We therefore aim to determine whether accurate imputation of context-specific connectivity query results is possible despite missing data.

The first step is imputing expression values for drug/cell combinations that lack experimental data. Since the early days of microarrays, there have been efforts to impute missing microarray expression values caused by array defects or hybridization issues (e.g., array scratches, localized manufacturing defects, reagent spatters). The naïve approach, averaging over expression values for a given gene in other samples in the data set, was quickly improved upon by more principled methods, including k-nearest neighbors and SVD (Troyanskaya *et al.*, 2001), local least squares optimization (Kim *et al.*, 2005), and Bayesian prediction (Oba *et al.*, 2003). Several methods made use of time series information when available (Troyanskaya *et al.*, 2001; Saha *et al.*, 2013; Bar-Joseph *et al.*, 2003). Collaborative filtering methods have even been applied to this problem (Saha *et al.*, 2016; Wang and Tseng, 2012).

However, nearly all of these approaches address a different problem – one in which there is a limited fraction (e.g. under 20%) of missing genes for a given sample, and in which the missing data points are not correlated across samples. A few approaches try to fit some characteristics of the random missing data, such as the observed histogram of the fraction of missing genes per sample (Oba *et al.*, 2003). Few of the time series methods (e,g, (Bar-Joseph *et al.*, 2003)) deal with imputing whole expression profiles.

Furthermore, to impute entirely missing expression profiles, one must incorporate additional domain information, such as data from nearby time points or functional relationships between genes’ expression patterns. In another example, transfer learning has been used to impute entire bulk RNA-sequencing profiles when methylation profiles for the same samples are available (Zhou *et al.*, 2020). Here, we use expression profiles of related drugs and cells.

Explicitly imputing connectivity for unassayed drugs and cells is thus a novel problem. Further, connectivity expression matrices can be very sparse, with well over half of the potential cell by drug matrix consisting of missing values. The novelty of our approach lies in using what data we do have about how specific drugs perform in other cells, and how various cells respond differently to other drugs, to infer missing data and then to assess efficacy specifically with respect to drug connectivity inference. The closest prior work we have seen is that of (Gottlieb *et al.*, 2017), which uses (a limited number of) expression values inferred from eQTL and patient cohort data to predict appropriate doses for warfarin. Such methods have great potential, but cannot be used without data on sequence variation.

## 2 METHODS

### 2.1 Overview of approach

To assess our ability to determine connectivity with missing data, we need a data set where we know the right answers. We create two such data sets by taking a complete subset of the LINCS connectivity data and a more realistic sparse subset in which approximately 75% of the drug/cell pairs are missing. We evaluate performance through five-fold cross-validation on each, as described below, to assess how well connectivity queries, performed with expression signatures characterizing each drug’s impact on cells, can produce the same results with missing data as with the full data. Specifically, given a drug-cell combination of interest, which we denote as drug *d_i_* and cell *c_j_*, and for which the expression profile is unavailable, we ask how well we can impute drug connectivity for *d_i_* and *c_j_*, and whether we can do so more accurately by taking cell type into account.

### 2.2 Connectivity mapping data and queries

Let *C* be a set of *c* cell types, and let *D* be a collection of *d* drugs, compounds, or cellular perturbations. Perturbations in LINCS include gene overexpression or knockdowns, but for the purposes of this study we consider primarily treatment of cells with named compounds or chemicals. We start with a matrix *M* of gene expression profiles for a common set of *n* genes in a subset of *D × C*. The matrix of cells and drugs is sparse, but the expression profiles are all-or-nothing; if we have expression data for some cell/drug combination, we have expression values for all *n* genes in the treated cell compared to the untreated cell. As in the paper introducing the L1000 connectivity data set (Subramanian *et al.*, 2017), these expression changes are represented by z-scores. Specifically, we use the published “Level 5” z-score data, which include z-scores of expression changes in drug-treated cells relative to controls, averaged over at least three replicates. When a cell / drug combination appears multiple times in the LINCS database, usually because that combination has been tested at different dosage levels or had expression profiles taken at different times after application, the expression profiles were combined into a consensus profile, using Stouffer’s method for combining Z-scores (Stouffer, 1949).

A *query signature* of a particular biological or disease state is defined to consist of two sets, one containing the *k* most up-regulated, and the other the *k* most down-regulated, genes in that state compared to a suitable control. So for example, a query signature for prostate cancer might consist of the 50 most up- and down-regulated genes in tumors from patients compared to normal prostate tissue.

A connectivity map query is performed using the query signature in the following way, as described in more detail in (Subramanian *et al.*, 2017). Given the query signature *q_u_* containing the *k* most upregulated and *q_d_* containing the *k* most downregulated genes for a given cell type and drug pair *c, d* and a reference expression profile *r*, we compute the weighted connectivity score (WTCS) as:

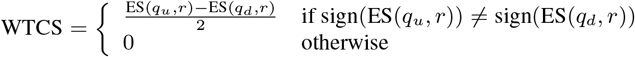

where ES(*q, r*) is the weighted Kolmogorov-Smirnov enrichment statistic (ES) described in (Subramanian *et al.*, 2005), and captures the enrichment of the set of genes *q* in profile *r*. WTCS ranges from −1 to 1. A score of 1 represents high positive connectivity, meaning that the drug’s effect on the given cell appears to be similar to that in the query signature, while a score of −1 represents high negative connectivity, or a drug/cell combination that up-regulates the down-regulated genes from the query signature and down-regulates its up-regulated genes.

Our data are derived from the LINCS data published in the Gene Expression Omnibus (GEO) (Edgar *et al.*, 2002) as GSE70138 and GSE92742. Together these contain 591,699 expression profiles for 98 cell types and 29,668 perturbagens spread over 189,173 unique cell/perturbagen combinations. The data were downloaded on Feb. 28, 2018.

For assessment purposes, we created two data matrices: a small, complete data set, and a larger, sparse data set. The complete data matrix contains 12 cell types: 8 cancer cell lines, A375, A549, HCC515, HEPG2, HT29, MCF7, PC3, and VCAP; HA1E, an immortalized normal kidney cell line; one stem cell line, ASC (normal adipose-derived stem cells), and two primary cell types: NPC (neural progenitor cells partially differentiated from iPSCs) and NEU (fully differentiated neurons). To create a complete and *interpretable* data matrix, we chose 450 drugs with “real” names (i.e., they are not just numbered compounds in development) and for which there is expression data for all 12 of the cell types above.

The sparse data matrix includes 80 cell types: 60 cancer cell lines, 6 immortalized normal cell lines, 4 stem cell lines, and 10 primary cell types. We selected a set of 1330 named drugs, again for interpretability, and expression data if available for each of the 80 cells listed. About 75% of the drug/cell combinations in this data set are missing, meaning that the compound was not assayed in that cell.

### 2.3 Data imputation methods

#### 2.3.1 Baseline methods

In the original connectivity map paper (Lamb *et al.*, 2006), connectivity scores were computed without consideration of cell type, essentially averaging across all cells. Given that the vast majority of these profiles were in a single cell line (MCF7), ignoring cellular context made sense. Even the current connectivity tool, using the much larger and more varied LINCS data set, reports averaged “summary” profiles (Subramanian *et al.*, 2017). This informs the idea behind our baseline imputation methods.

##### Tissue-agnostic

A good baseline prediction of drug *d_i_*’s performance in cell *c_j_* might be simply to look at what drug *d_i_* does to a cell, regardless of what type of cell it is. Assume that we have expression profiles for drug *d_i_* on other cell types. By taking the median gene expression profile over of that drug all the other cells for which we do have data (the highlighted row in Figure 1a), we arrive at a prediction of what drug *d_i_* “usually does” to a cell. We call this the *tissue agnostic* imputation method, and compare other results to this.

**Fig. 1.**
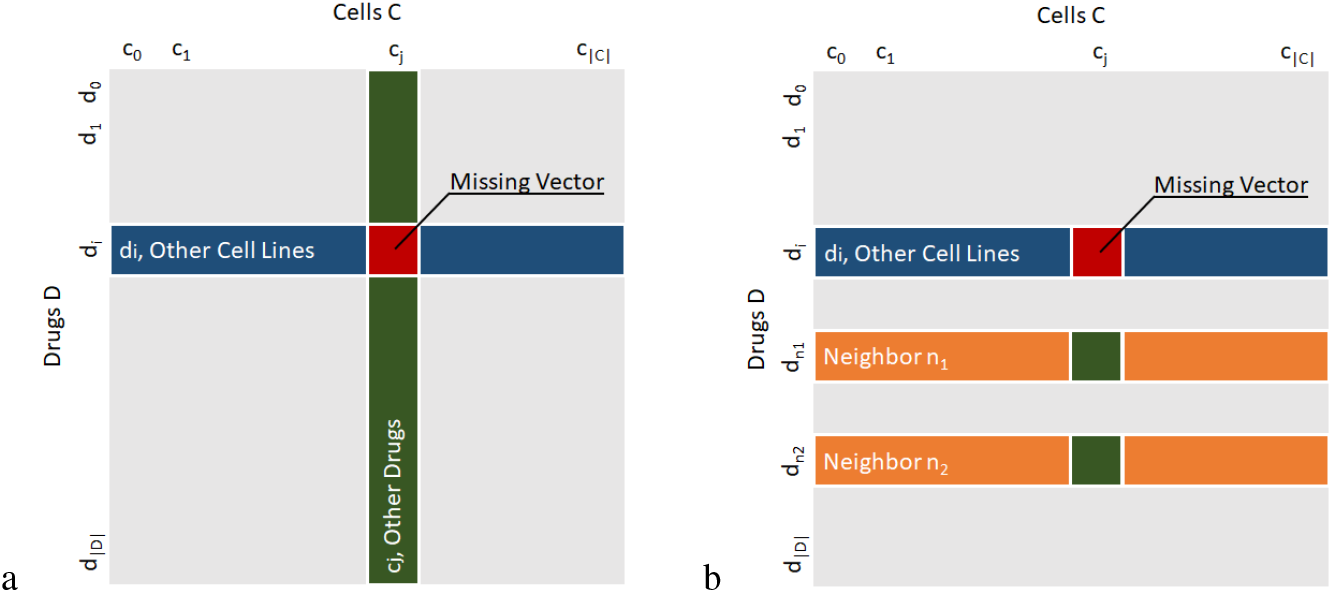
Methods overview. a) In tissue-agnostic imputation, the median of the expression profiles (vectors) for drug *d_i_* on cells other than *c_j_* (in blue) is used to predict the unknown expression profile for drug *d_i_* on cell *c_j_*. The two-way average is the average of two sets of expression profiles: the median of the expression profiles for drug *d_i_* on cells other than *c_j_* (in blue) and the median of the expression profiles for drugs other than *d_i_* on cell *c_j_* (in green). b) In the neighborhood approach to collaborative filtering imputation, neighbors are drugs whose expression profiles on cells other than *c_j_* (in orange) are most similar to *d_i_*’s profile on cells other than *c_j_* (in blue). In this example with *k* = 2 neighbors, the expression profile for cell *c_j_* on neighbor drugs *d_n_*_1_ and *d_n_*_2_, shown in green, are used to impute the profile for drug *d_i_* on cell *c_j_*.

##### Two-way average

We might additionally want to include tissue-specific information in a very straightforward way. We can accomplish this by averaging the tissue-agnostic prediction for drug *d_i_* across all other cells with the analogous “drug-agnostic” prediction for *c_j_* (the highlighted column in Figure 1a) that tells us how expression of *c_j_* after perturbation typically differs from expression in other cells. We refer to this as *two-way average* prediction.

#### 2.3.2 Collaborative filtering

Collaborative filtering is an approach used in recommender systems to impute missing rating values and thereby recommend new products to users based both on information from similar users and from other items that user has rated. Calculations are typically based on sparse databases with *m* users and *n* items containing those users’ ratings for ≤ *n* of those items (Su and Khoshgoftaar, 2009). These ratings can represent any kind of relationship between users and items. In applications such as movie or purchase recommendations, the ratings might be represented by integers in the range [1,5]. But in other applications, ratings might be real numbers or categorical variables. This approach is of particular interest for imputation of connectivity data because of the sparsity of the database.

##### Neighborhood approach

One approach to collaborative filtering is to rely on the closest neighbors of a particular sample as a model. An average, weighted by similarity of the neighbors’ ratings, approximates the rating for the sample of interest. A critical change was necessary to use this approach with our data, because each “rating” in the connectivity matrix is actually a vector of gene expression values. To avoid loss of information, we view the gene expression values as a multi-part rating of the same item. In the user/movie-rating metaphor, these values would represent a rating that perhaps specified an overall rating, as well as ratings of the movie’s acting, cinematography, and soundtrack. If two users rate a movie similarly but one rates the cinematography poorly and other rates it highly, these users might actually be more different than they first appear.

From the data matrix *M* defined in section 2.2, we compute a “ratings” matrix *R* by mean-centering the row of a drug’s “rating vectors” (expression values) for all cells. Specifically, let *μ* be the mean of all non-missing values in the row, and then replace each existing value *v* by (*v – μ*) and each missing value by (0 – *μ*).

We then calculate similarities by taking the cosine similarity between all pairs of rows in *R*, yielding a symmetric matrix.

For each missing expression profile (vector), we then predict that profile by finding the top *x* “neighbors” of the row containing the missing value. We used *k* = 50 neighbors for the complete 12×450 data set and *k* = 120 neighbors for the sparse 80×1330 data set; both are around ten percent of the number of rows in the matrix. (We tuned the fraction of neighbors parameter on a smaller and older data set consisting of 200 non-overlapping drugs and 6 cells; we did not find the results to be highly sensitive to changes in *k*. Additionally, we replicated our results for the 80×1330 data set, below, with *k* increased or decreased by a factor of two, and again confirmed that we see minimal effects on the connectivity results.)

We compute the predicted profile *P_ij_* for drug *d_i_* and cell *c_j_*, as

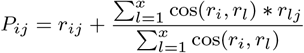

where cos refers to the cosine distance between two ratings vectors, *r_ij_* refers to the ratings vector in row *i* and column *j*, *r_i_* is the average value of all values present in row *i*, and *r_lj_* is the value corresponding to the current drug-cell-gene combination in the neighbor row *l*.

##### Matrix decomposition

The goal of this approach is to account for latent subclasses within the drugs or cells. We used the FunkSVD package, which applies stochastic gradient descent SVD optimization to build an approximation of an original input matrix *M*. This specific methodology was shown to be particularly effective in predicting Netflix movie ratings (Bennett and Lanning, 2007). The FunkSVD method does not require a complete matrix to run, and effectively “overlooks” missing or unknown ratings (Funk, 2006). Note that this is not the case for a simple SVD decomposition, which would require some initial “guess” for the missing entries (Sarwar *et al.*, 2000). Therefore, we worked directly with the raw values in *M*. We still consider *M* to be a multi-part ratings matrix, although it is not mean-centered like *R*.

FunkSVD decomposes the matrix into component matrices *U* and *V* with singular values folded into those matrices. The parameters of this function determine the output rank of the approximation. For our purposes we used rank parameter *k* = 55. (As with the nearest neighbors approach, we tuned the rank parameter *k* on the smaller, older data and we evaluated the 80×1330 data set with 2x and 1/2x *k*; again, changes in *k* did not greatly impact the results.) A lower-rank approximation can then be obtained by reconstructing the matrix with *M* = *UV′*. We then predicted the values of missing data per row with the approximation *r*_new_ = *uV′* where *u* is a row in *M* containing an unknown multi-part rating.

### 2.4 Evaluation

#### 2.4.1 Cross-validation

To assess our performance, we need data on which we know the right answers. For each of the complete connectivity matrix and the sparse matrix described in section 2.2, we created cross-validation data sets in the following way.

Each cell / drug combination is randomly and independently assigned to one of five folds. We then verify that the candidate set of fold assignments has no fold where more than 75 percent of the cells for a given drug, or 75 percent of the drugs for a given cell, are assigned to that fold, ensuring that any method would be able to produce an imputed expression profile for any missing cell/drug combination. If this requirement was violated, fold assignments were completely regenerated until the requirement was met.

For each fold, a given method is provided *only* the z-score normalized gene expression profiles for cell / drug combinations not in the fold, and must impute the expression profiles for cell / drug combinations that are in the fold. For the sparse data set, folds consist of true profiles only for evaluation; missing data is assigned a fold value of zero and predictions for such values do not factor into our accuracy assessments. Over all five folds, a given method will produce a single imputed profile for each cell / drug combination. We then compare the imputed profile to the true profile for that cell / drug combination, using the various scoring metrics described below.

To assess variance due to the randomness in this cross-validation procedure, we created five independent instances (“runs”) of cross validation data sets for both the complete and sparse matrices. We summarize our results across those five runs.

#### 2.4.2 Scoring connectivity prediction

Once we have a predicted expression profile for a drug-cell combination, we must assess how accurate our prediction is through some comparison to the known true expression profile. Since our aim is to infer connectivity for cell-drug pairs for which we lack data, an informative evaluation method would be to compare the connectivity results from the withheld true data with those obtained using the imputed data.

There are two parts of a list of drugs returned by a connectivity query that are of interest. Drugs with the most positive connectivity scores are drugs that replicate the query signature; these are sometimes used to identify similar drugs, or to find compounds that might cause a similar change to that of the query expression profile as an adverse event. Drugs with the most negative connectivity scores are those that reverse the observed query signature; these are candidate therapeutics for an observed disease signature. Accordingly, we want to assess how well the most-positive, or most-negative, connectivity results from the imputed data match those from the withheld true data.

To do this, we use the Weighted Spearman Rank Correlation measure, defined by (Shieh *et al.*, 2000). We use a weight function defined as

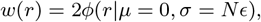

where *r* is the result rank, *ϕ* is the normal distribution of the form *ϕ*(*x|μ, σ*), and *ϵ* is a parameter with value between 0 and 1 controlling the “aggressiveness” of the curve.

In this work, we chose *ϵ* = 0.01, which applies weights of significant magnitude to approximately the top 20 results, after which weights diminish towards zero. We tested the effect of doubling or halving *ϵ* but saw minimal changes in the results.

### 2.5 Detecting drugs with similar functions

#### 2.5.1 Do predictions connect to true signatures of similar drugs

Beyond comparison to true profiles, we can evaluate our performance more directly by assessing the connectivity of a drug’s expression profile to other drugs within the same perturbagen class. A perturbagen class (PCL) is a group of drugs that share a mechanism of action or target genes. PCLs are defined by published literature and further refined by connectivity analysis to yield 171 classes, as described in more detail in (Subramanian *et al.*, 2017). Actual PCL membership used in this manuscript was obtained through personal communication with the authors.

For a given compound, a query signature reflecting expression changes caused by treatment of cells with that compound is expected to return strong positive connectivity between that drug and others in the same PCLs. To evaluate a method, we can construct a query from an imputed signature and determine if drugs in the same PCL set are over-represented towards the top of the returned list, expressing strong positive connectivity.

To do this, we used the imputed expression profile for each d rug, *d*, to create a query signature with the top and bottom 50 genes (the number recommended by (Subramanian *et al.*, 2017)). We then use that signature to query the remaining drugs in the true data set, returning a list of weighted enrichment scores for each drug, ordered from most to least positively connected to drug *d*.

We adapt gene set enrichment analysis tools (Subramanian *et al.*, 2005; Planet, 2019) to function as a drug set enrichment analysis, evaluating the distribution of drugs within a PCL set in a connectivity result. Ideally the drugs other than the one that produced the query signature should be among the most connected compounds.

For this analysis, we require a minimum of three drugs per PCL. After removing drug classes with no more than 2 drugs *in the chosen query matrix*, there are 132 drugs of the 450 in the complete data set that are in the remaining perturbagen classes. In the sparse data set, 334 of the 1330 drugs are in PCLs.

An enrichment score (ES) is calculated as described in 2.2, weighted by connectivity scores from the query results. In this context, an ES captures enrichment of a PCL set in the connectivity result of a drug that is a member of that set. The normalized enrichment score (NES) is calculated to account for the varying set sizes. Significance of enrichment is determined by comparing the observed NES to the NES of 1,000 random permutations of the connectivity result. The fraction of the absolute value of the permuted NES greater than, and thus stronger, than the absolute value of the observed NES yields the normalized p-value (Mootha, 2003). (For negative connectivity scores this becomes the fraction of random NES less than the observed NES.)

For each drug/cell combination the imputed signature is queried against the true matrix. The distribution of the normalized enrichment scores calculated for each drug, pcl, and cell combination is compared across methods, with the expectation that the NES should be strongly positive if strong connectivity is correctly detected.

#### 2.5.2 Finding similar unassayed drugs

With known drug classes, we can also assess how well we impute connectivity query results *to* drug/cell combinations that were not experimentally evaluated. This allows us to compare efficacy of connectivity imputation across cells and drug classes. Missing data is predicted from the true expression profiles using neighborhood collaborative filtering, chosen because it was the best approach overall. Experimentally evaluated signatures are replaced with the average of their imputed profiles from each of the five runs to generate a completely imputed matrix from the sparse data set. We also created another completely imputed matrix using the tissue agnostic method for comparison. Each signature is queried against the complete matrix, with strong positive connectivity between drugs in the same PCL correctly identified if the NES score is positive and significant (meaning the normalized p value is less than 0.05). We measure the percentage of drugs significantly enriched in their PCL for every PCL/cell combination to assess our ability to impute connectivity query results even in the absence of experimental data, as well as to estimate the amount of data needed.

## 3 RESULTS

### 3.1 Using cell-specific data improves imputation

To assess accuracy, we performed connectivity queries with query signatures containing the most dysregulated genes for each drug in the treated cell line compared to control. Specifically, the query signatures consist of the 50 most up- and down-regulated genes for a given drug. Positive connectivity scores identify the drugs that induce expression changes most similar to those of the query drug; negative connectivity scores identify drugs whose effects on a cell reverse the effect of the query drug.

For the complete data set, Figures 2 and 3 show the average weighted Spearman correlations (over all drugs) between the true and predicted connectivity results (a) and the percent changes for each compared to the tissue agnostic method (b). Corresponding figures for the sparse matrix can be found in the supplemental data. These are organized both by cell type and by the percentage of drugs in a given cell that have been profiled experimentally.

**Fig. 2.**
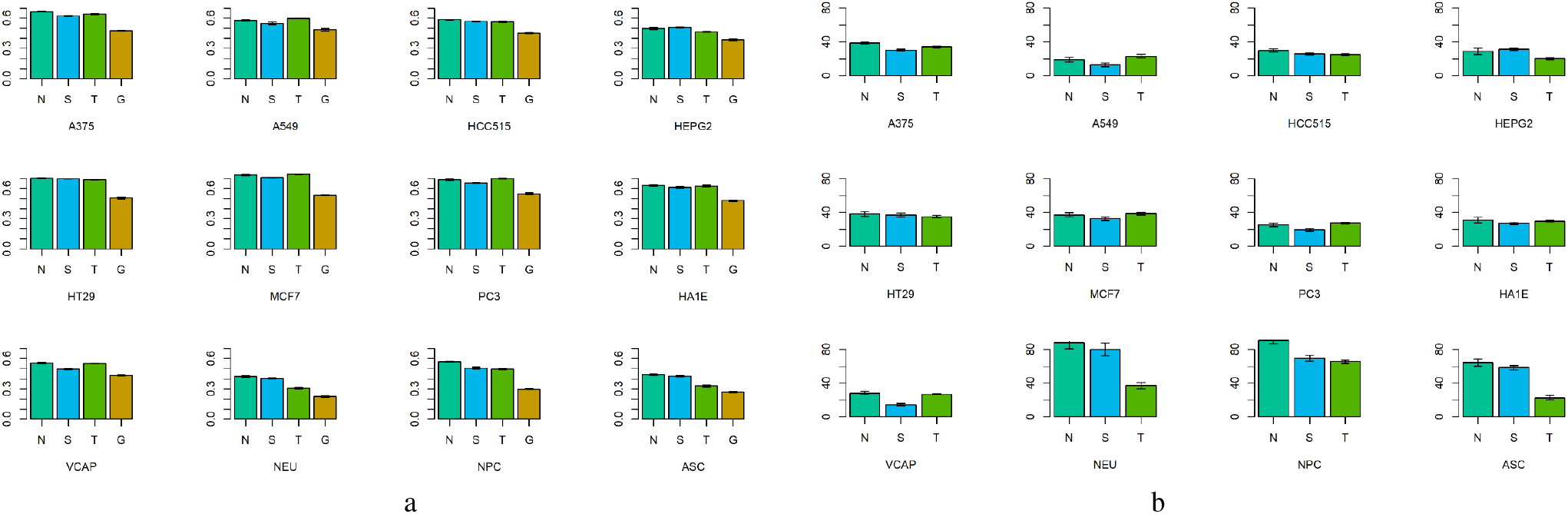
a) Negative weighted connectivity correlation across all genes and drugs, for each of the 12 cell lines in the complete matrix. Methods are denoted by single-letter labels: N : neighborhood collaborative approach; S : SVD; T : two-way average; G : tissue-aGnostic (baseline method). Error bars show variation across cross validation runs. b) Percent change in negative weighted connectivity correlation compared to the tissue-agnostic method.

**Fig. 3.**
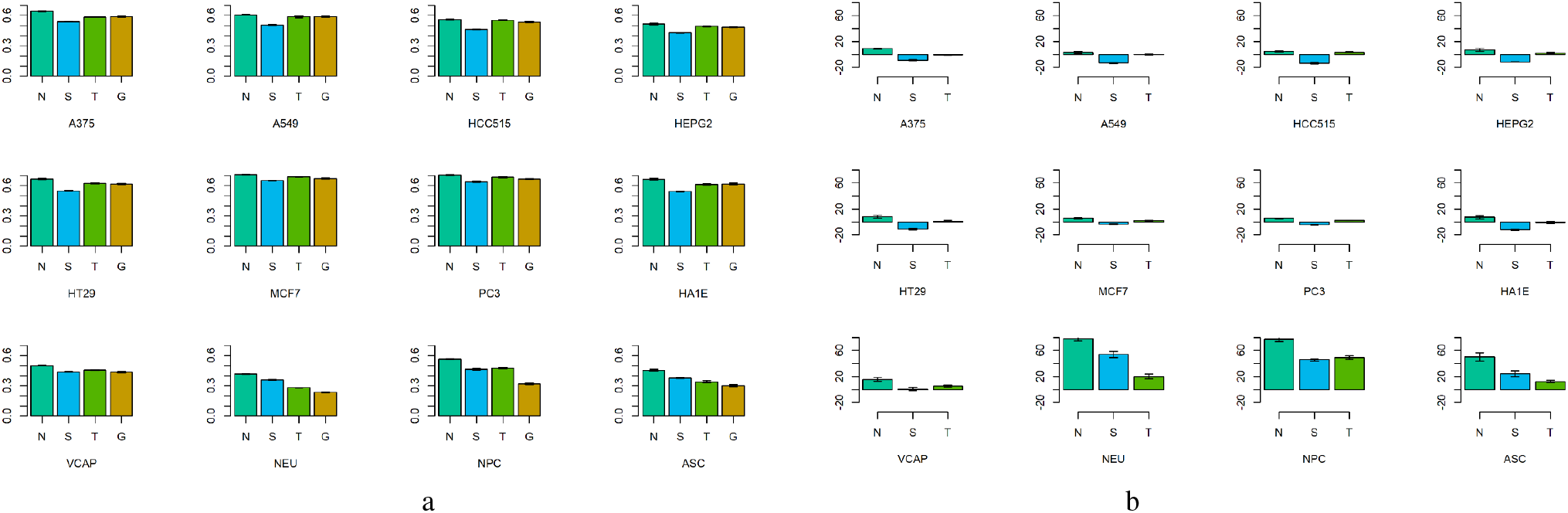
a) Positive weighted connectivity correlation across all genes and drugs, for each of the 12 cell types in the complete matrix. Error bars show variation across cross validation runs. Labels are the same as in Figure 2. b) Percent change in positive weighted connectivity correlation compared to the tissue-agnostic method.

For the complete matrix, negative connectivity, which is the most commonly envisioned use case of the connectivity map, shows robust improvement in prediction under almost all methods, with neighborhood collaborative filtering either best or essentially tied for best with the two-way average in all cells. The neighborhood method’s improvements are especially large for the primary cell types, where it notably out-performs the two-way average. But even in the cancer cell lines, an improvement of 20-40% over the tissue-agnostic baseline is seen with the neighborhood method.

Positive imputation results show little improvement over baseline for all but the primary and stem cell types in the complete matrix. Indeed, SVD methods are actually worse than tissue-agnostic imputation here. But the neighborhood collaborative filtering and two-way average approaches, both of which use cell specific information, are at least as good as the tissue-agnostic approach for the immortalized cells. For the primary and stem cells, the neighborhood approach still shows a substantial improvement, with two-way average and SVD vying for second place. Note that the tissue-agnostic weighted correlation for NEU cells is the lowest (0.24) in this data set, with the other primary cell, NPC, in the low 0.30s, and most immortalized cell lines showing a weighted correlation between 0.5 and 0.67. Thus, improving on these much higher scores may be harder.

Overall these results demonstrate that cell type is an important aspect of connectivity mapping. We conclude that incorporating cell-specific information improves connectivity outcomes in all cells, but it does so most dramatically in cells that are neither malignant nor immortalized.

In the sparse matrix, there is improvement over baseline for both positive and negative connectivity correlations across all classes of cells. The nearest neighbors approach continues to outperform all methods for cells with enough data, with SVD and two-way average improving over tissue agnostic as well.

However, this trend begins to break down for cells where the percentage of drugs assayed is low. For primary cell types, tissue-specific methods are more powerful in all cells with at least 41% of the 1330 drugs assayed. In cells containing data for 5% or less of the compounds, i.e. where there is not much cell-specific data to work with, the tissue agnostic method is sometimes a more reliable approach, and error bars reflecting the varying cross validation folds become notably larger.

This holds true for cancer cell lines as well, where tissue-specific methods and particularly the neighborhood approach exceed tissue agnostic imputation for cells with at least 48% of the compounds assayed, but it becomes much less robust for cells in which fewer than 13% of the compounds have data, and here again there are large error bars. Immortalized and stem cells, though there are far fewer of them with reasonable amounts of data, appear to follow a similar pattern.

### 3.2 Downsampling reveals amount of data needed

To determine how performance varies as a function of the number of compounds assayed, filling the gap between 41% and 13% identified in the previous section, we selected a few cells from each category of cells with more than 75% of the compounds assayed and downsampled the number of drugs. Compounds were randomly removed from folds until just 50%, 40%, 30%, 20% or 10% of the drugs remained for a given cell line. Signatures were imputed once more and negative connectivity results were compared to the true results. This was repeated across all five instances of cross validation data sets.

Figure 4 shows the negative weighted connectivity correlation for each cell type averaged across all genes, drugs, and runs, plotted against percent of compounds used for imputation. We found that in all cells analyzed, performance didn’t suffer notably so long as at least 20% of the drugs had data (e.g. we assayed approximately 260 of the 1330 compounds). Additionally, so long as data were available for at least 10% of the drugs, all methods performed better than tissue agnostic regardless of cell type, with nearest neighbors having the best performance or, in a few cases, being a close second.

These early findings suggest that for the larger, sparser data set, incorporating cellular context improves positive and negative connection detection over all cells types that have data for at least 130 drugs. Further work will be needed to determine more precisely how the distribution of compounds sampled affects predictive performance.

**Fig. 4.**
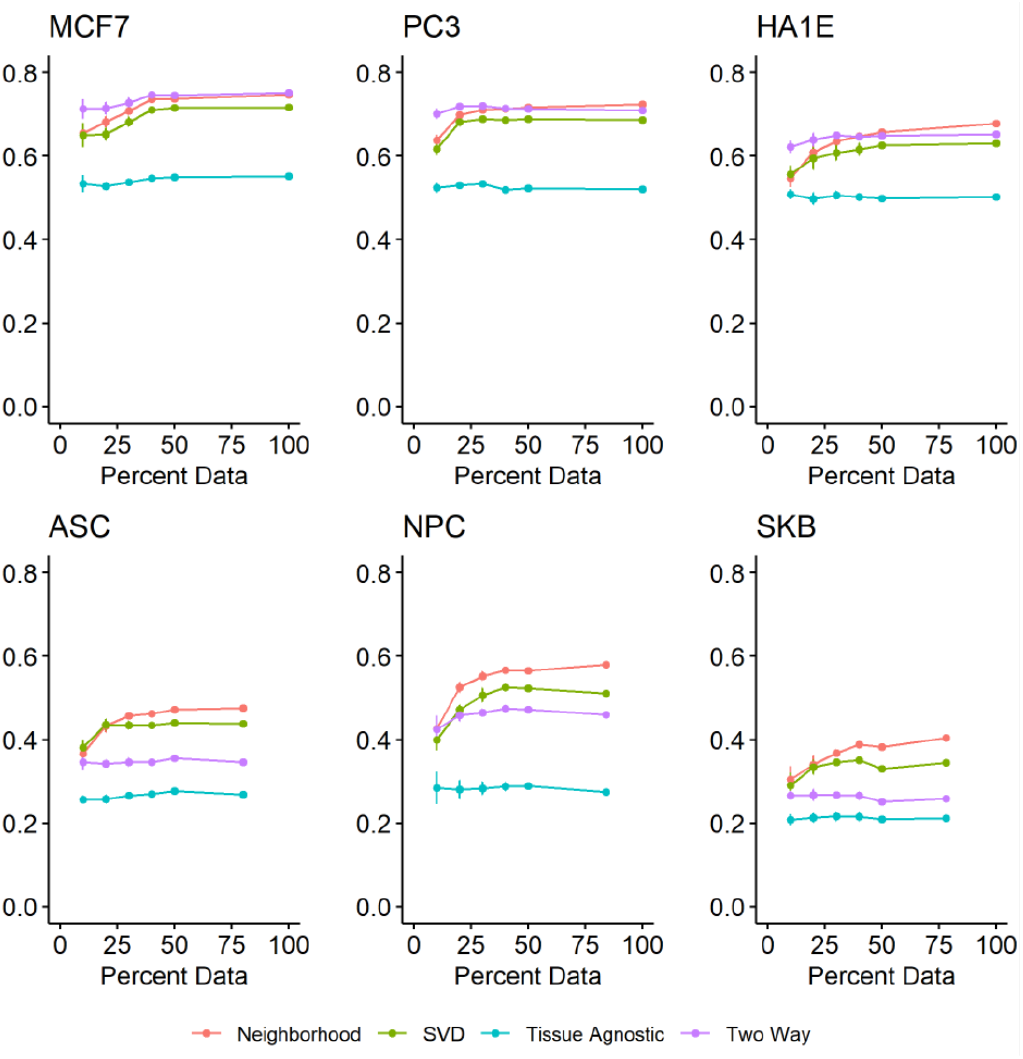
Negative weighted connectivity correlation for the named cells across all genes and drugs (y-axis), as a function of the percentage of the 1330 compounds whose data was available for imputation. Maximum x-values on each plot reflect the actual amount of data for that cell; all other points reflect downsampling. Error bars show the variation across cross-validation runs; most are smaller than the data markers.

### 3.3 Imputed profiles detect related compounds

Figure 5 shows the distribution of NES scores between a drug and its corresponding Perturbagen Class (PCL) in a given cell for the complete matrix (a) and sparse matrix (b), using the method described in section 2.5.1. In this context, a positive NES score means the connectivity results for drug *d* has drugs within the same PCL set clustered towards the top of the returned list, thus strong connectivity is accurately detected. For randomized lists NES scores are expected to be 0; if the imputed profiles successfully replicate the signature of drugs in the same PCLs, then we would expect the greatest density of NES scores to be well above zero.

**Fig. 5.**
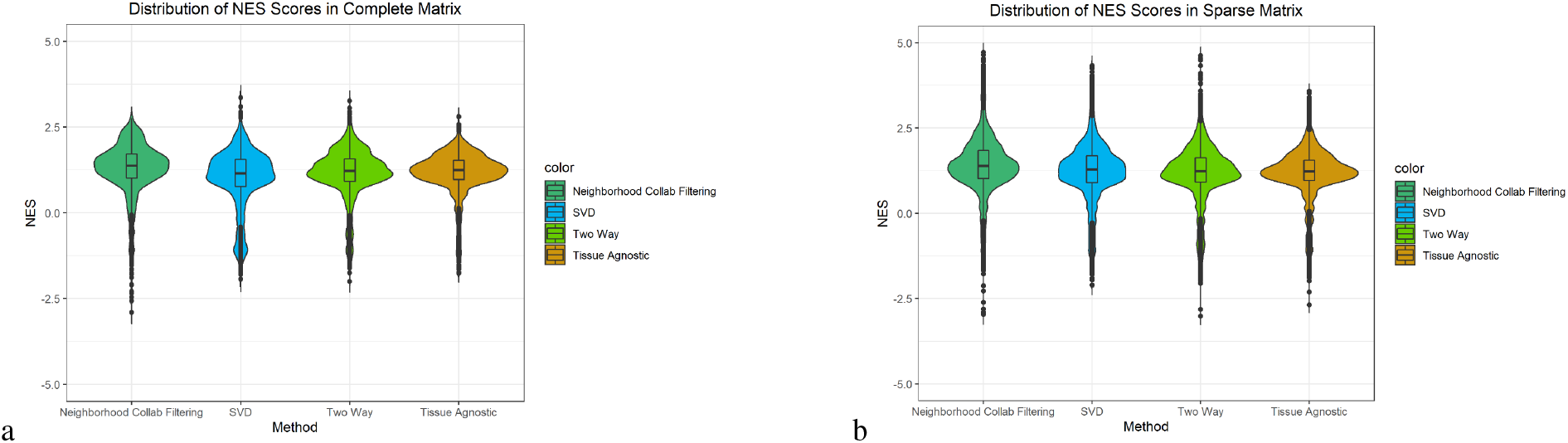
a) Violin plot showing distribution of normalized enrichment scores of drugs from a given drug set imputed from the complete matrix. b) Violin plot showing distribution of normalized enrichment scores of drugs from a given drug set imputed from the sparse matrix.

For all methods in both the complete and sparse matrix, the median of the NES scores is greater than 1.0, with the majority of NES scores clustered well above zero. Neighborhood collaborative filtering has a slightly higher median than all other methods and greater NES values overall in the sparse matrix, proving to be the most robust and consistent method to be evaluated.

### 3.4 Predicting connectivity of unassayed drugs

Given the performance of the neighborhood collaborative filtering method in Section 3.1, we generated a complete version of the sparse matrix (where 75% of the cells were missing) using only imputed data, as described in Section 2.5.2. We then assessed our ability to find related drugs within the same PCL using imputed query profiles against this completely imputed matrix.

Figure 6a shows the percentage of drugs enriched for their PCL set using this complete matrix. Just primary cells are shown, with full plots containing all 80 cell types in the supplemental data, organized by cell type (cancer, immortalized, stem and primary) and ordered by percentage of drugs profiled in each cell and PCL size.

**Fig. 6.**
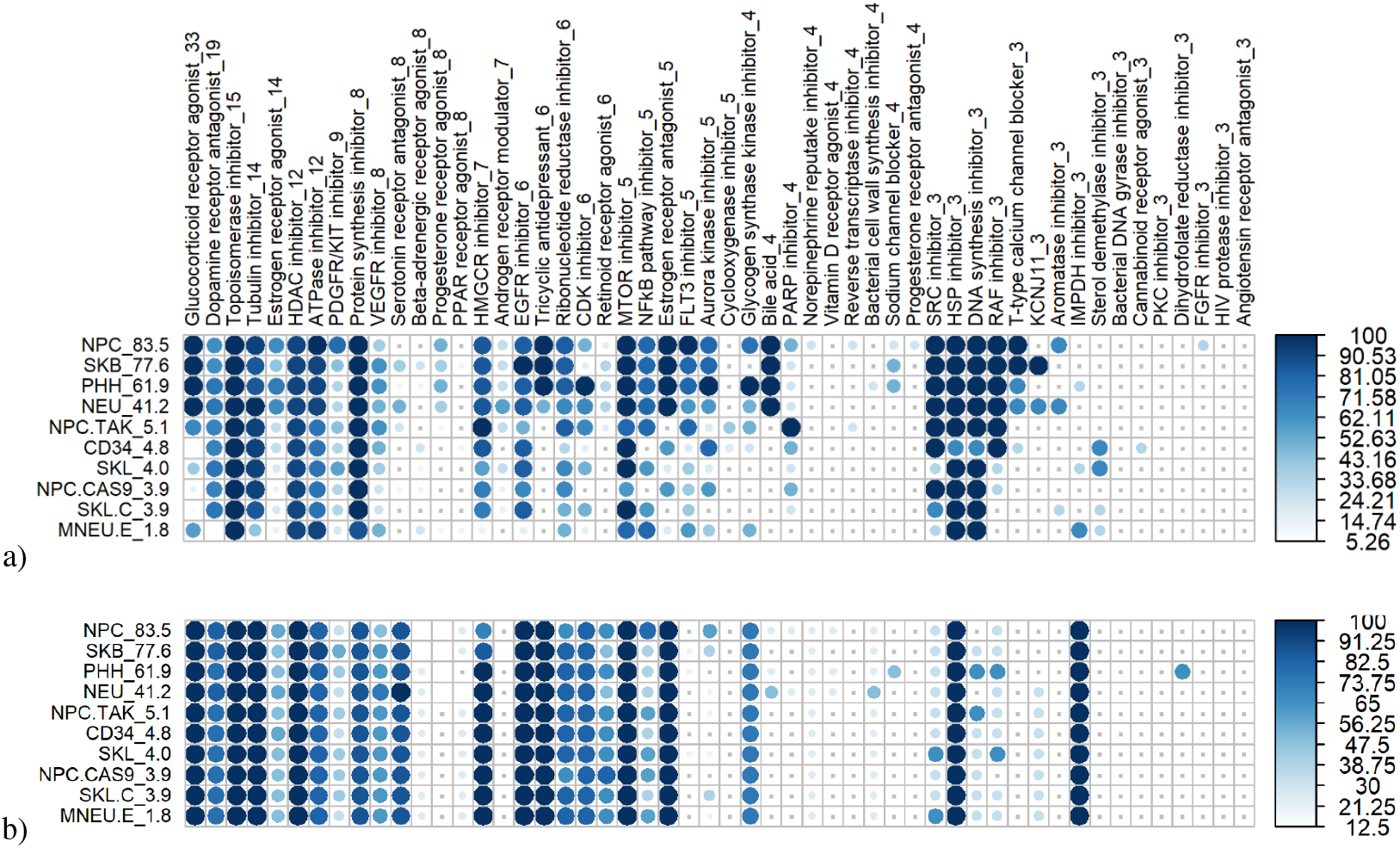
a) Percent of drugs correctly expressing strong connectivity to their drug class using the fully imputed sparse matrix by the neighborhood approach, shown only for primary cells. The number of drugs in each PCL set is included in the PCL name, and the percentage of drugs assayed in each cell type is included in the cell name. The darker the shade of blue, the higher percentage of drugs with statistically significant NES scores. Grey dots represent PCL/cell combinations in which there were no statistically significant NES scores. b) Percent of drugs correctly expressing strong connectivity to their drug class using the fully imputed sparse matrix by the tissue agnostic approach, shown only for primary cells. The darker the shade of blue, the higher percentage of drugs with statistically significant NES scores. Grey dots represent PCL/cell combinations in which there were no statistically significant NES scores. Plots were generated using the corrplot package (Wei and Simko, 2017).

Of the drug/cell/PCL combinations, 53% have normalized p-values less than 0.05, meaning that in the majority of drug and cell combinations evaluated, connectivity query results are correctly imputed by the neighborhood collaborative filtering method, given a sparse matrix of experimental data. The approach was best at accurately detecting strong positive connectivity in most of the drug sets in NPC, SKB, PHH and NEU cells, which had data for over 41% of the compounds. In cells containing data for 5% or less of the compounds, the percentage of significant NES scores becomes less reliable, consistent with the results noted above for positive and negative weighted connectivity correlations. This is true across all cell types assessed, with cells containing less than 5% of the 1330 drugs assayed showing a lower percentage of statistically significant NES scores.

Figure 6b shows the same plot for data imputed by the tissue-agnostic method, allowing us to identify classes of drugs in which imputation is better using cell-type-specific or tissue-agnostic methods. We notice that for tissue-agnostic imputation, the amount of data for the individual cell predictably doesn’t have much affect on accuracy; the bottom cells (with much less data) look more or less the same as those on top.

Similarly, drug classes with the most drugs tend be well imputed across the board, suggesting the likelihood that in such cases the “neighborhood” is well populated by related drugs. Drug classes for which imputation is better with collaborative filtering include aurora kinase inhibitors, FLT3 inhibitors, and a number of steroid/hormone classes (progesterone receptor agonists, androgen receptor modulators). Classes where tissue-agnostic methods were better included retinoid receptor agonists and serotonin receptor antagonists. (These trends are visible in the primary cells in Figure 6 but may be clearer in the supplemental figure showing all cells.)

We conclude that positive connections between drugs of the same class can be indeed be imputed for missing data. To do this well, more than 5% of the compounds from the 1330 assessed need to be experimentally evaluated for robust imputation, but again further work will be needed to determine more precisely the impact the number and distribution of compounds assayed has on accurate connectivity prediction.

### 3.5 Examples of predicted connectivity in disease

To illustrate use cases of our collaborative-filtering predictions, we identified differentially expressed genes in a previously-published microarray study that compared the bulk transcriptomes of postmortem hippocampus samples between subjects with schizophrenia and age- and sex-matched controls ((Lanz *et al.*, 2019); GSE53987). We applied the fifty most up- and down-regulated genes as the “query signature” against a hybrid version of the sparse matrix, in which missing data was replaced by our cell-specific predictions. Our goal was to query both assayed and imputed data to identify compounds that might reverse this neuronal transcriptional signature despite not having been assayed in neurons (denoted NEU) in the LINCS data set.

Most of the top compounds reversing the schizophrenia signature had indeed already been directly profiled in neurons, so validating their links to schizophrenia simply supports the idea of connectivity mapping in non-cancer cells. Still, as an illustration, the top two such compounds were pirfenidone, an anti-fibrotic that has shown neurological effects in multiple sclerosis and chronic pain (Walker and Margolin, 2001; Peng *et al.*, 2020), and remoxipride, an atypical antipsychotic indicated for schizophrenia, but whose use since 2003 has been limited due to toxicity concerns (Nadal, 2001).

The most connected compounds *missing* from the sparse matrix, and that thus would not have been discovered without our predictive approach, include theophylline and bosentan. Theophylline is a naturally-occurring phosphodiesterase- and HDAC-inhibitor typically used in asthma but known to have neurological effects. It has been suggested for potential therapeutic use in both schizophrenia (Zagorska *et al.*, 2018) and Alzheimer’s disease (Fernando *et al.*, 2017). Bosentan is an endothelin receptor antantagonist whose targets, EDNRA and EDNRB, regulate migration of neural progenitor cells (Nishikawa *et al.*, 2011); mouse mutations of these genes lead to abnormal neurological morphology and function (www.informatics.jax.org). These surprising yet plausible connections, along with more expected ones (e.g., citalopram, a selective serotonin reuptake inhibitor inexplicably not profiled in neurons), were discoverable only through predictive methods.

We next queried the same database with the signature of differentially expressed genes from a microarray study that compared left ventricular myocardial samples from ischemic cardiomyopathy patients to controls ((Kong *et al.*, 2010) GSE16499). There are no myocardial (heart muscle) cells in the LINCS data sets we downloaded, but there are primary skeletal muscle cells (SKL), although with data for only 4% of compounds their predictive power may be limited. Nonetheless, the most connected predicted compounds for the cardiomyopathy signature are givinostat, shown to have therapeutic efficacy in a mouse model of heart failure (Jeong *et al.*, 2018), and the ATP-ase inhibitor brefeldin-a, which affects cholesterol transport (Verghese *et al.*, 2008), a process important in cardiovascular disease. The one compound more significantly connected in muscle cells and actually assayed in the SKL cells was vorinostat (also known as suberoylanilide hydroxamic acid or SAHA), which has also shown benefit in a number of models of cardiac dysfunction (Chelladurai *et al.*, 2021).

## 4 DISCUSSION

We have demonstrated that context-specific connectivity data can be used to infer missing data in a connectivity data matrix. This has applications for the full LINCS data set, in which many cell types have data for just a few tens of compounds, and even beyond. We have also determined that for our sparse matrix covering 1330 drugs assessed, cell-specific information does not always improve imputation of connectivity when fewer than 5% of the drugs have been profiled in the cell. Downsampling the number of compounds for cell lines with the most available data suggested that at least 20% of the data is necessary for reliable cell-specific imputation. An important direction for future work is to better understand the impact of the distribution of compounds needed for robust prediction across sparse matrices of different sizes and compositions.

Regarding method comparisons for imputation, the neighborhood collaborative filtering approach has been most successful in this study. That said, there is the possibility of improving SVD by using higher-order methods that better capture the three-dimensional structure of the data. Such approaches require considerable parameter tuning on a distinct data set. Beyond that, there is room for additional improvement using different machine learning methods.

We have further demonstrated that both negative and positive connectivity can be inferred from a complete imputed matrix generated from the sparse data set. Drugs that are members of PCLs with at least 12 compounds of the 1330 drugs analyzed are enriched for other members of their PCL set across all cell types with enough data. Of all the drug/cell/PCL combinations tested, 53% have a statistically significant NES score. This suggests the imputed expression profiles replicate the imputed query signatures of drugs in the same class, and the complete imputed matrix can be used for finding drugs with the same mechanism of action or, alternatively, drugs that reverse disease states.

Indeed, queries based on actual expression data from patients were able to identify compounds that were not tested in those cells and would not have been discovered without these predictive approaches. Of the four compounds highlighted as new discoveries in our example disease queries, only givinostat would have been among the top 10% of compounds if querying the “summary” profile, an average of just the assayed cells. The others are only discoverable through cell-specific imputation.

Finally, we have demonstrated that considering cell type is critical for accurate connectivity mapping. This was apparent in prior work using multiple different breast cancer cell lines (Niepel *et al.*, 2017), but becomes even more important across more varied contexts. Both for precision cancer medicine purposes and for use in applications beyond oncology, taking context into account is therefore essential.

## ACKNOWLEDGEMENTS

We thank Ted Natoli and Aravind Subramanian for their help in data interpretation, for sharing the PCL lists, and for feedback on earlier versions of this work. We also thank members of the Tufts BCB research group, especially Dan Meyer, for helpful suggestions.

## REFERENCES

Bar-Joseph, Z., Gerber, G. K., Gifford, D. K., Jaakkola, T. S., and Simon, I. (2003). Continuous representations of time-series gene expression data. J. Comput. Biol., 10(3), 341–356.

Bennett, J. and Lanning, S. (2007). The netflix prize.

Chelladurai, P., Boucherat, O., Stenmark, K., Kracht, M., Seeger, W., Bauer, U. M., Bonnet, S., and Pullamsetti, S. S. (2021). Targeting histone acetylation in pulmonary hypertension and right ventricular hypertrophy. Br J Pharmacol, 178(1), 54–71.

Edgar, R., Domrachev, M., and Lash, A. E. (2002). Gene Expression Omnibus: NCBI gene expression and hybridization array data repository. Nucleic Acids Res., 30(1), 207–210.

Fernando, W. M. A. D. B., Somaratne, G., Goozee, K. G., Williams, S., Singh, H., and Martins, R. N. (2017). Diabetes and Alzheimer,s Disease: Can Tea Phytochemicals Play a Role in Prevention? J Alzheimers Dis, 59(2), 481–501.

Funk, B. W. S. (2006). Netflix update: Try this at home. https://sifter.org/simon/journal/20061211.html.

Gottlieb, A., Daneshjou, R., DeGorter, M., Bourgeois, S., Svensson, P. J., Wadelius, M., Deloukas, P., Montgomery, S. B., and Altman, R. B. (2017). Cohort-specific imputation of gene expression improves prediction of warfarin dose for African Americans. Genome Med, 9, 98.

Jeong, M. Y., Lin, Y. H., Wennersten, S. A., Demos-Davies, K. M., Cavasin, M. A., Mahaffey, J. H., Monzani, V., Saripalli, C., Mascagni, P., Reece, T. B., Ambardekar, A. V., Granzier, H. L., Dinarello, C. A., and McKinsey, T. A. (2018). Histone deacetylase activity governs diastolic dysfunction through a nongenomic mechanism. Sci Transl Med, 10(427).

Kim, H., Golub, G. H., and Park, H. (2005). Missing value estimation for DNA microarray gene expression data: Local least squares imputation. Bioinformatics, 21, 187–198.

Kong, S. W., Hu, Y. W., Ho, J. W., Ikeda, S., Polster, S., John, R., Hall, J. L., Bisping, E., Pieske, B., dos Remedios, C. G., and Pu, W. T. (2010). Heart failure-associated changes in RNA splicing of sarcomere genes. Circ Cardiovasc Genet, 3(2), 138–146.

Lamb, J., Crawford, E. D., Peck, D., Modell, J. W., Blat, I. C., Wrobel, M. J., Lerner, J., Brunet, J. P., Subramanian, A., Ross, K. N., Reich, M., Hieronymus, H., Wei, G., Armstrong, S. A., Haggarty, S. J., Clemons, P. A., Wei, R., Carr, S. A., Lander, E. S., and Golub, T. R. (2006). The Connectivity Map: using gene-expression signatures to connect small molecules, genes, and disease. Science, 313(5795), 1929–1935.

Lanz, T. A., Reinhart, V., Sheehan, M. J., Rizzo, S. J. S., Bove, S. E., James, L. C., Volfson, D., Lewis, D. A., and Kleiman, R. J. (2019). Postmortem transcriptional profiling reveals widespread increase in inflammation in schizophrenia. Transl Psychiatry, 9(1), 151.

Mootha, V., L. C. E. K. (2003). Pgc-1-responsive genes involved in oxidative phosphorylation are coordinately downregulated in human diabetes. Nature Genetics, 34, 267–273.

Nadal, R. (2001). Pharmacology of the atypical antipsychotic remoxipride, a dopamine D2 receptor antagonist. CNS Drug Rev, 7(3), 265–282.

Niepel, M., Hafner, M., Duan, Q., Wang, Z., Paull, E. O., Chung, M., Lu, X., Stuart, J. M., Golub, T. R., Subramanian, A., Ma’ayan, A., and Sorger, P. K. (2017). Common and cell-type specific responses to anti-cancer drugs revealed by high throughput transcript profiling. Nat Commun, 8(1), 1186.

Nishikawa, K., Ayukawa, K., Hara, Y., Wada, K., and Aoki, S. (2011). Endothelin/endothelin-B receptor signals regulate ventricle-directed interkinetic nuclear migration of cerebral cortical neural progenitors. Neurochem Int, 58(3), 261–272.

Oba, S., Sato, M., Takemasa, I., Monden, M., K., M., and Ishii, S. (2003). A Bayesian missing value estimation method for gene expression profile data. Bioinformatics, 19(16), 2088–2096.

Peng, X., Guo, H., Chen, J., Wang, J., and Huang, J. (2020). The effect of pirfenidone on rat chronic prostatitis/chronic pelvic pain syndrome and its mechanisms. Prostate, 80(12), 917–925.

Planet, E. (2019). phenoTest: Tools to test association between gene expression and phenotype in a way that is efficient, structured, fast and scalable. We also provide tools to do GSEA (Gene set enrichment analysis) and copy number variation. R package version 1.34.0.

Saha, S., Dey, K., Dasgupta, R., Ghose, A., and Mullick, K. (2013). Missing value estimation in DNA microarrays using B-splines. Journal of Medical and Bioengineering, 2(2), 88=92.

Saha, S., Ghosh, A., and Nath Dey, K. (2016). An improved fuzzy based approach to impute missing values in DNA microarray gene expression data with collaborative filtering. In International Conference on Advances in Computing, Communications and Informatics, ICACCI –16, pages 911–916. IEEE.

Sarwar, B., Karypis, G., Konstan, J., and Riedl, J. (2000). Application of dimensionality reduction in recommender system - a case study. Technical report, University of Minnesota.

Shieh, G. S., Bai, Z., and Tsai, W.-Y. (2000). Rank tests for independence – with a weighted contamination alternative. Statistica Sinica, 10(2), 577–593.

Stouffer, S. (1949). A study of attitudes. Sci. Am., 14(5), 11–15.

Su, X. and Khoshgoftaar, T. M. (2009). A survey of collaborative filtering techniques. Adv. in Artif. Intell., pages 4:2–4:2.

Subramanian, A. et al. (2017). A Next Generation Connectivity Map: L1000 Platform and the First 1,000,000 Profiles. Cell, 171(6), 1437–1452.

Subramanian, A., Tamayo, P., Mootha, V. K., Mukherjee, S., Ebert, B. L., Gillette, M. A., Paulovich, A., Pomeroy, S. L., Golub, T. R., Lander, E. S., et al. (2005). Gene set enrichment analysis: a knowledge-based approach for interpreting genome-wide expression profiles. Proceedings of the National Academy of Sciences, 102(43), 15545–15550.

Troyanskaya, O., Cantor, M., Sherlock, G., Brown, P., Hastie, T., Tibshirani, R., Botstein, D., and Altman, R. B. (2001). Missing value estimation methods for dna microarrays. Bioinformatics, 17(6), 520–5.

Verghese, P. B., Arrese, E. L., Howard, A. D., and Soulages, J. L. (2008). Brefeldin A inhibits cholesterol efflux without affecting the rate of cellular uptake and re-secretion of apolipoprotein A-I in adipocytes. Arch Biochem Biophys, 478(2), 161–166.

Walker, J. E. and Margolin, S. B. (2001). Pirfenidone for chronic progressive multiple sclerosis. Mult Scler, 7(5), 305–312.

Wang, B.-W. and Tseng, V. S. (2012). Improving missing-value estimation in microarray data with collaborative filtering based on rough-set theory. International Journal of Innovative Computing, Information and Control, 8.3(B), 2157–2172.

Wei, G., Twomey, D., Lamb, J., Schlis, K., Agarwal, J., Stam, R., Opferman, J., Sallan, S., den Boer, M., Pieters, R., Golub, T., and Armstrong, S. (2006). Gene expression-based chemical genomics identifies rapamycin as a modulator of MCL1 and glucocorticoid resistance. Cancer Cell, 10(4), 331–42.

Wei, T. and Simko, V. (2017). R package –corrplot–: Visualization of a Correlation Matrix. (Version 0.84).

Zagorska, A., Partyka, A., Bucki, A., Gawalskax, A., Czopek, A., and Pawlowski, M. (2018). Phosphodiesterase 10 Inhibitors - Novel Perspectives for Psychiatric and Neurodegenerative Drug Discovery. Curr Med Chem, 25(29), 3455–3481.

Zhang, C., Ryu, Y., Chen, T., Hall, C., Webster, D., and Kang, M. (2012). Synergistic activity of rapamycin and dexamethasone in vitro and in vivo in acute lymphoblastic leukemia via cell-cycle arrest and apoptosis. Leuk Res, 36(3), 342–9.

Zhou, X., Chai, H., Zhao, H., Luo, C., and Yang, Y. (2020). Imputing missing rna-sequencing data from dna methylation by using a transfer learning-based neural network. Gigascience, 9(7), giaa076.

