## Supplemental Table and Figures for "Cell-specific imputation of drug connectivity mapping with incomplete data"

May 11, 2021

Contents:

1. All 80 cells in sparse matrix organized by cell type (Table 1).
2. Positive connectivity results for sparse 80x1330 matrix (Fig. 1).
3. Percent change in positive connectivity results compared to tissue agnostics: sparse 80x1330 matrix (Fig. 2).
4. Negative connectivity results for sparse 80x1330 matrix (Fig. 3)
5. Percent change in negative connectivity results compared to tissue agnostics: sparse 80x1330 matrix (Fig. 4).
6. Percent of drugs enriched for PCL set in full neighborhood matrix: all cell types (Fig. 5).
7. Percent of drugs enriched for PCL set in full tissue agnostic matrix: all cell types (Fig. 6).
8. Comparison between various $\epsilon$ settings for the weighted correlation score between connectivity queries (Fig. 7).
9. Comparison between halving and doubling parameter k: positive connectivity (Fig. 8).
10. Comparison between halving and doubling parameter k: negative connectivity (Fig. 9).

| **Cancer** | **Primary** |
|---|---|
| MCF7, HT29, A375, PC3, A549, HCC515 , | NPC |
| VCAP, HEPG2, HELA, YAPC, U937, LOVO, | SKB |
| SNUC4, SKMEL1, RMUGS, HCC15, HEC108, CORL23, | PHH |
| A673, NCIH596, TYKNU, SW948, SW620, SNU1040, | NEU |
| SNGM, SKMEL28, OV7, RKO, NCIH508, H1299, | NPC.TAK |
| AGS, EFO27, SW480, HCT116, JHUEM2, MDST8, | CD34 |
| NCIH2073, COV644, DV90, RMGI, SKLU1, HT115, | SKL |
| WSUDLCL2, PL21, NCIH1836, NCIH1694, SNUC5, CL34, | NPC.CAS9 |
| THP1, SKM1, T3M10, NOMO1, BT20, HS578T, | SKL.C |
| MDAMB231, SKBR3, HUH7, JURKAT, LNCAP, HL60 | MNEU.E |
| **Immortalized** | **Stem Cell** |
| HA1E | ASC |
| HME1 | FIBRNPC |
| MCF10A | ASC.C |
| HUVEC | HUES3 |
| NKDBA | |
| HEK293T | |

Table 1: Cells Types and Cells in Sparse Data Set.

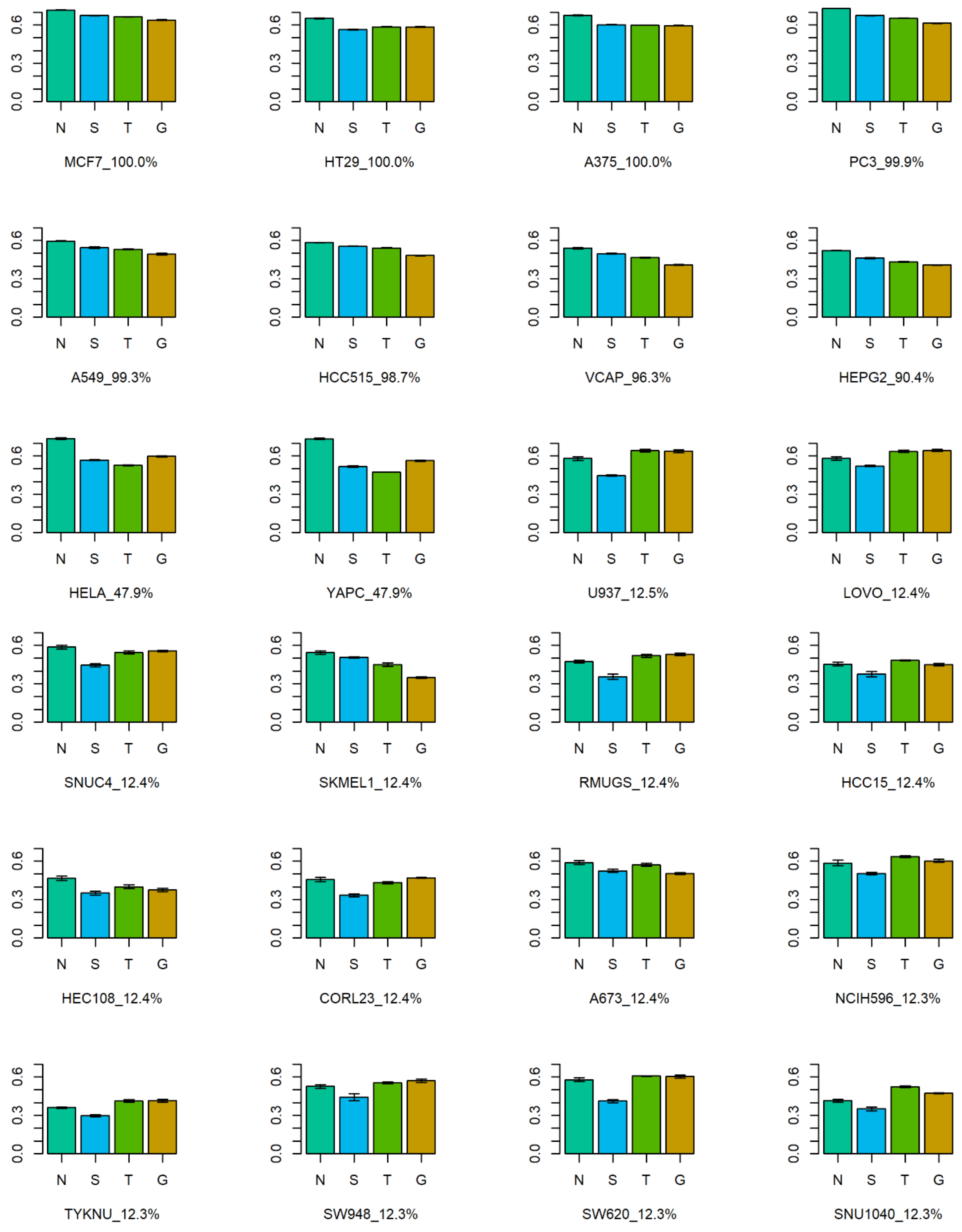
MCF7_100.0%
HT29_100.0%
A375_100.0%
PC3_99.9%
A549_99.3%
HCC515_98.7%
VCAP_96.3%
HEPG2_90.4%
HELA_47.9%
YAPC_47.9%
U937_12.5%
LOVO_12.4%
SNUC4_12.4%
SKMEL1_12.4%
RMUGS_12.4%
HCC15_12.4%
HEC108_12.4%
CORL23_12.4%
A673_12.4%
NCIH596_12.3%
TYKNU_12.3%
SW948_12.3%
SW620_12.3%
SNU1040_12.3%
N S T G
0.0 0.3 0.6

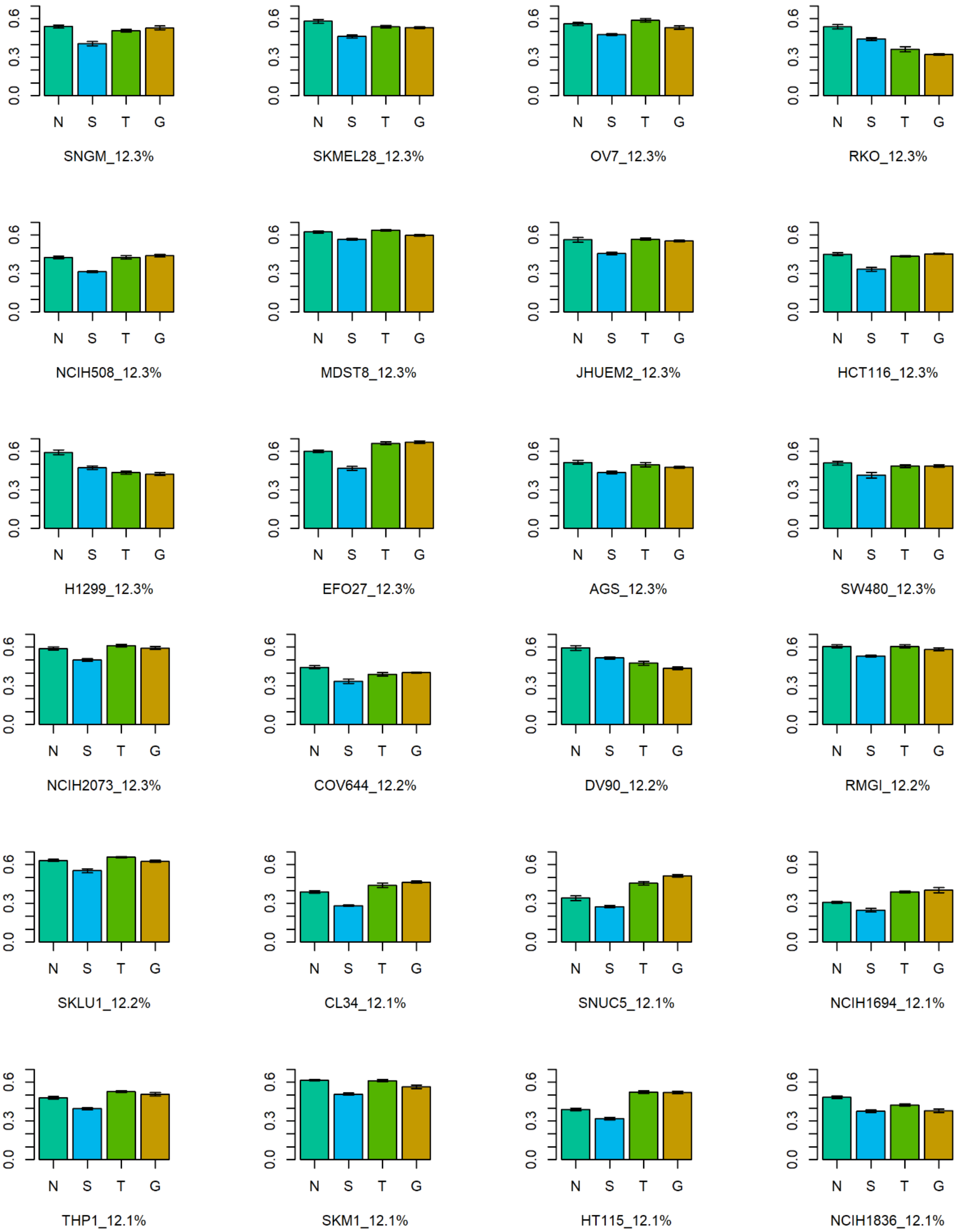
SNGM_12.3%
SKMEL28_12.3%
OV7_12.3%
RKO_12.3%
NCIH508_12.3%
MDST8_12.3%
JHUEM2_12.3%
HCT116_12.3%
H1299_12.3%
EFO27_12.3%
AGS_12.3%
SW480_12.3%
NCIH2073_12.3%
COV644_12.2%
DV90_12.2%
RMGI_12.2%
SKLU1_12.2%
CL34_12.1%
SNUC5_12.1%
NCIH1694_12.1%
THP1_12.1%
SKM1_12.1%
HT115_12.1%
NCIH1836_12.1%
N S T G
0.0 0.3 0.6

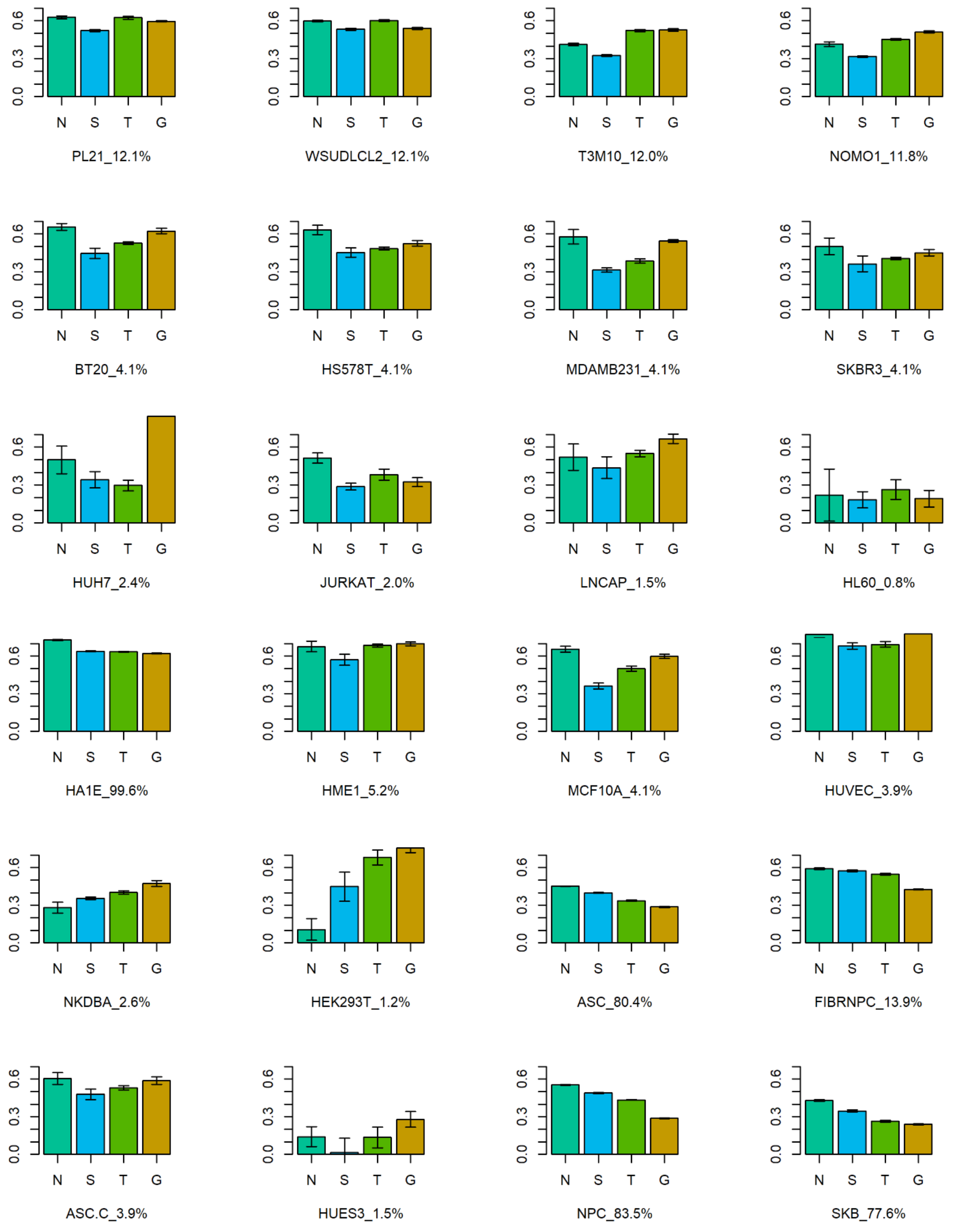
0.0
0.3
0.6
N S T G
PL21_12.1%
WSUDLCL2_12.1%
T3M10_12.0%
NOMO1_11.8%
BT20_4.1%
HS578T_4.1%
MDAMB231_4.1%
SKBR3_4.1%
HUH7_2.4%
JURKAT_2.0%
LNCAP_1.5%
HL60_0.8%
HA1E_99.6%
HME1_5.2%
MCF10A_4.1%
HUVEC_3.9%
NKDBA_2.6%
HEK293T_1.2%
ASC_80.4%
FIBRNPC_13.9%
ASC.C_3.9%
HUES3_1.5%
NPC_83.5%
SKB_77.6%

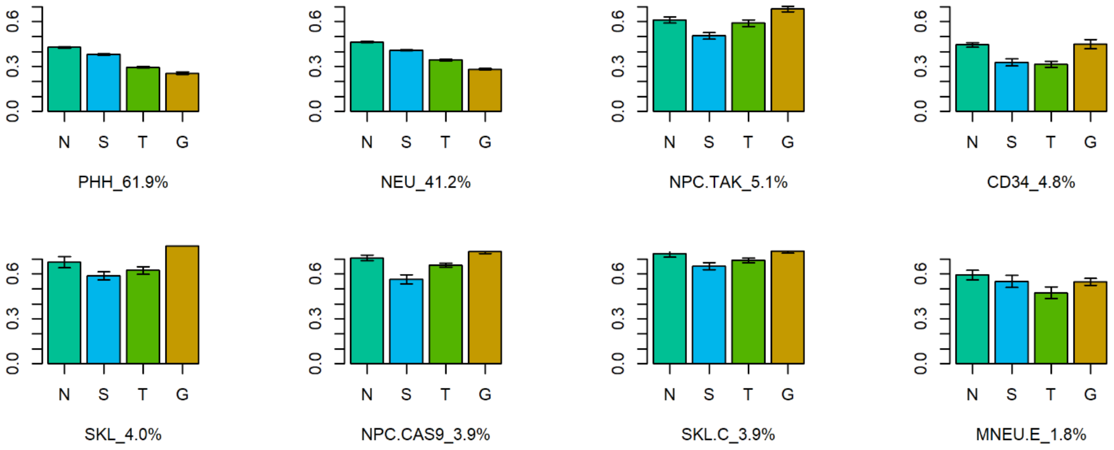


Figure 1: Positive weighted connectivity correlation across all genes and drugs, for each of the 80 cell lines in the sparse matrix. Methods are denoted by single-letter labels: N : neighborhood collaborative approach; S : SVD; T : two-way average; G : tissue-aGnostic (baseline method). Organized by cell type (cancer, immortalized, stem and primary) and ordered by percentage of drugs profiled in each cell. Error bars show variation across cross validation runs.

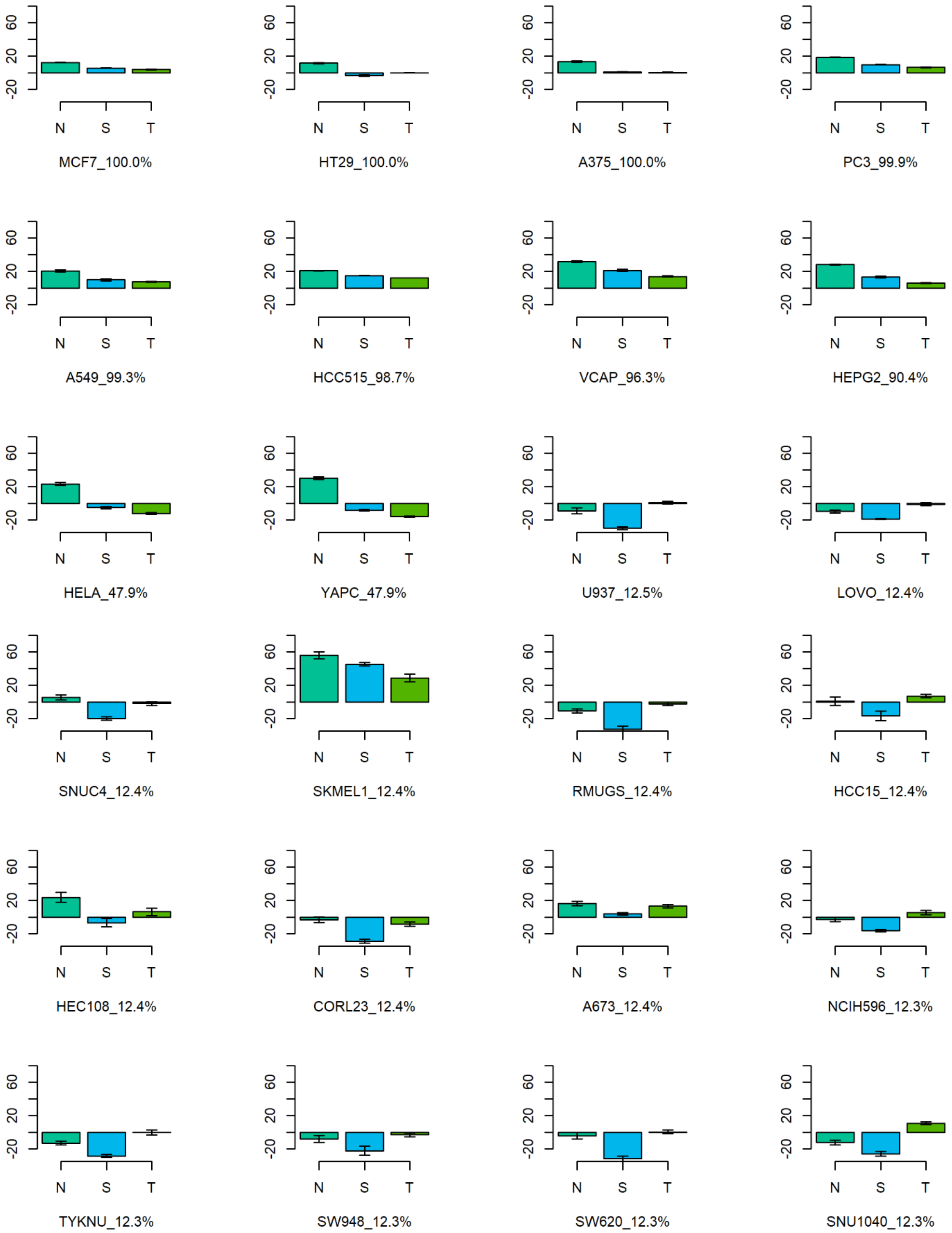
MCF7_100.0%
HT29_100.0%
A375_100.0%
PC3_99.9%
A549_99.3%
HCC515_98.7%
VCAP_96.3%
HEPG2_90.4%
HELA_47.9%
YAPC_47.9%
U937_12.5%
LOVO_12.4%
SNUC4_12.4%
SKMEL1_12.4%
RMUGS_12.4%
HCC15_12.4%
HEC108_12.4%
CORL23_12.4%
A673_12.4%
NCIH596_12.3%
TYKNU_12.3%
SW948_12.3%
SW620_12.3%
SNU1040_12.3%

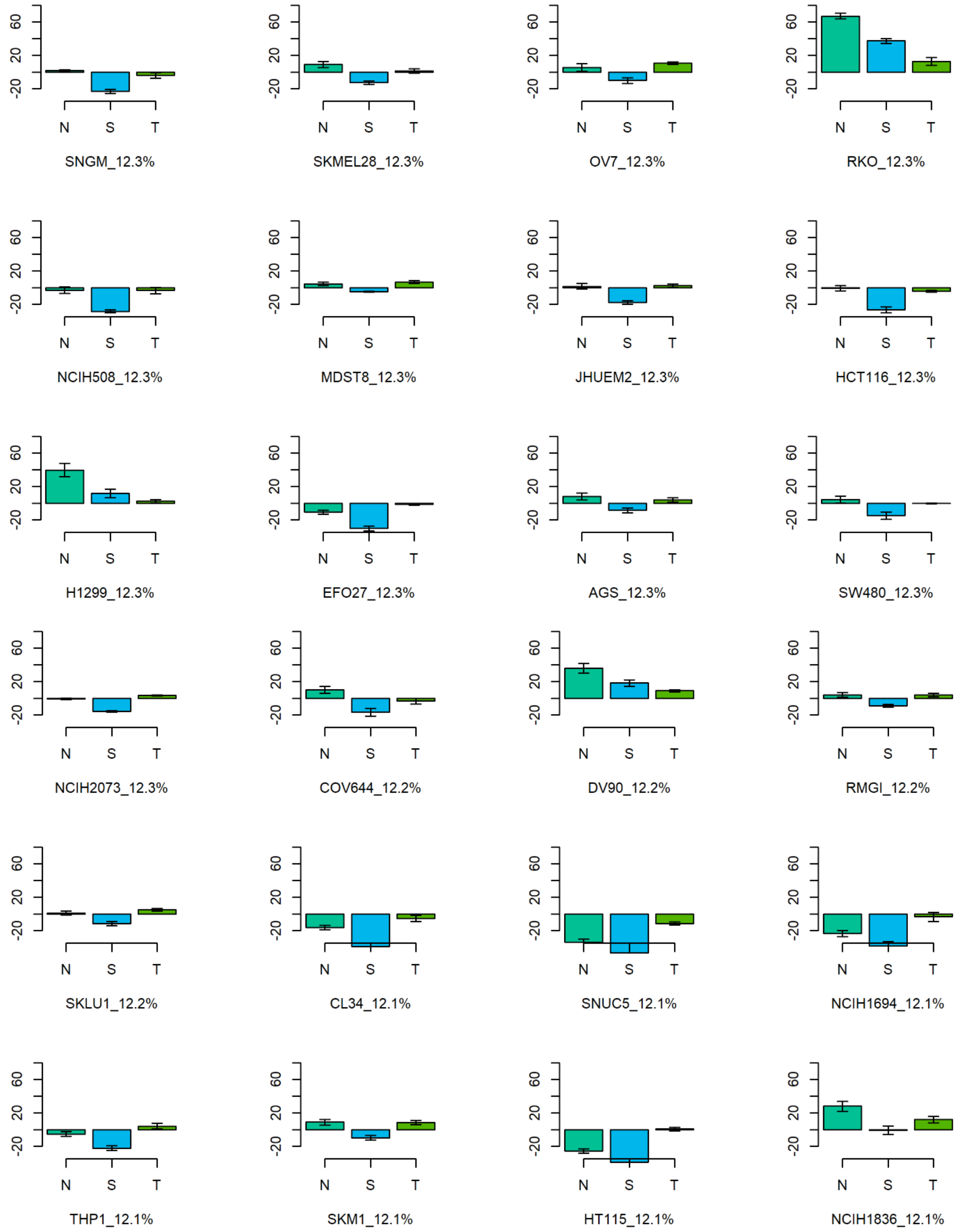
SNGM_12.3%
SKMEL28_12.3%
OV7_12.3%
RKO_12.3%
NCIH508_12.3%
MDST8_12.3%
JHUEM2_12.3%
HCT116_12.3%
H1299_12.3%
EFO27_12.3%
AGS_12.3%
SW480_12.3%
NCIH2073_12.3%
COV644_12.2%
DV90_12.2%
RMGI_12.2%
SKLU1_12.2%
CL34_12.1%
SNUC5_12.1%
NCIH1694_12.1%
THP1_12.1%
SKM1_12.1%
HT115_12.1%
NCIH1836_12.1%
N
S
T
60
20
-20

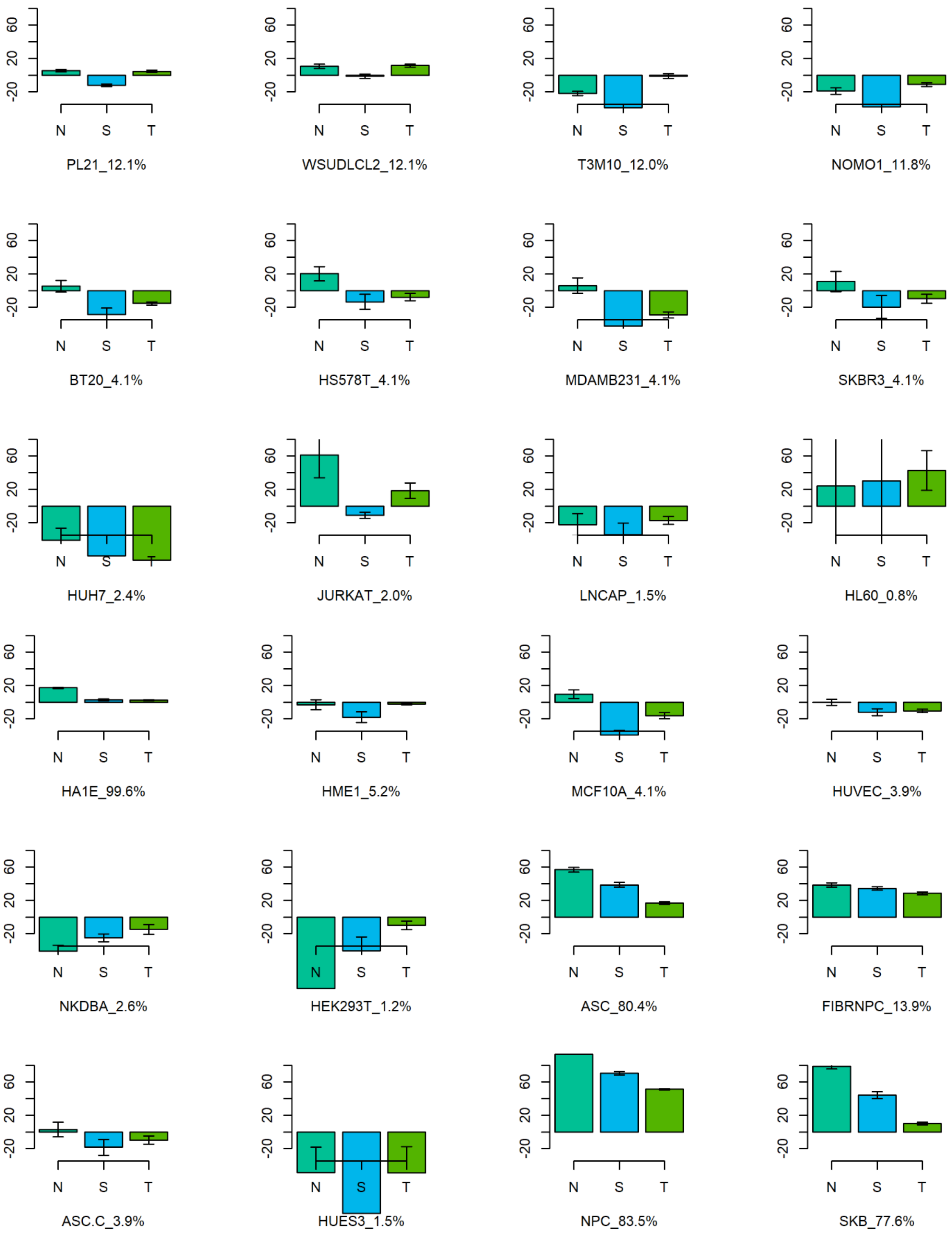
PL21_12.1%
WSUDLCL2_12.1%
T3M10_12.0%
NOMO1_11.8%
BT20_4.1%
HS578T_4.1%
MDAMB231_4.1%
SKBR3_4.1%
HUH7_2.4%
JURKAT_2.0%
LNCAP_1.5%
HL60_0.8%
HA1E_99.6%
HME1_5.2%
MCF10A_4.1%
HUVEC_3.9%
NKDBA_2.6%
HEK293T_1.2%
ASC_80.4%
FIBRNPC_13.9%
ASC.C_3.9%
HUES3_1.5%
NPC_83.5%
SKB_77.6%
N
S
T
60
20
-20

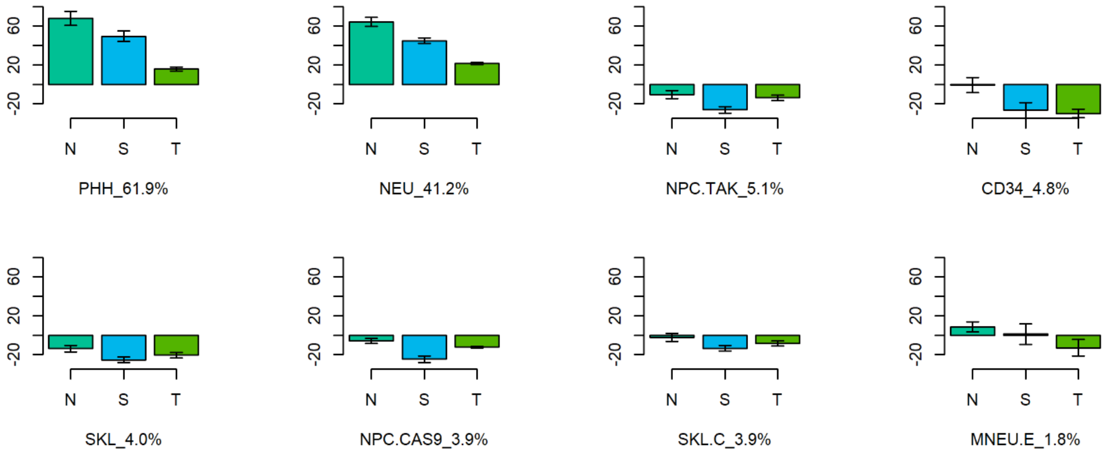


Figure 2: Percent change in positive weighted connectivity correlation compared to the tissue-agnostic method. Methods are denoted by single-letter labels: N : neighborhood collaborative approach; S : SVD; T : two-way average; G : tissue-aGnostic (baseline method). Organized by cell type (cancer, immortalized, stem and primary) and ordered by percentage of drugs profiled in each cell. Error bars show variation across cross validation runs.

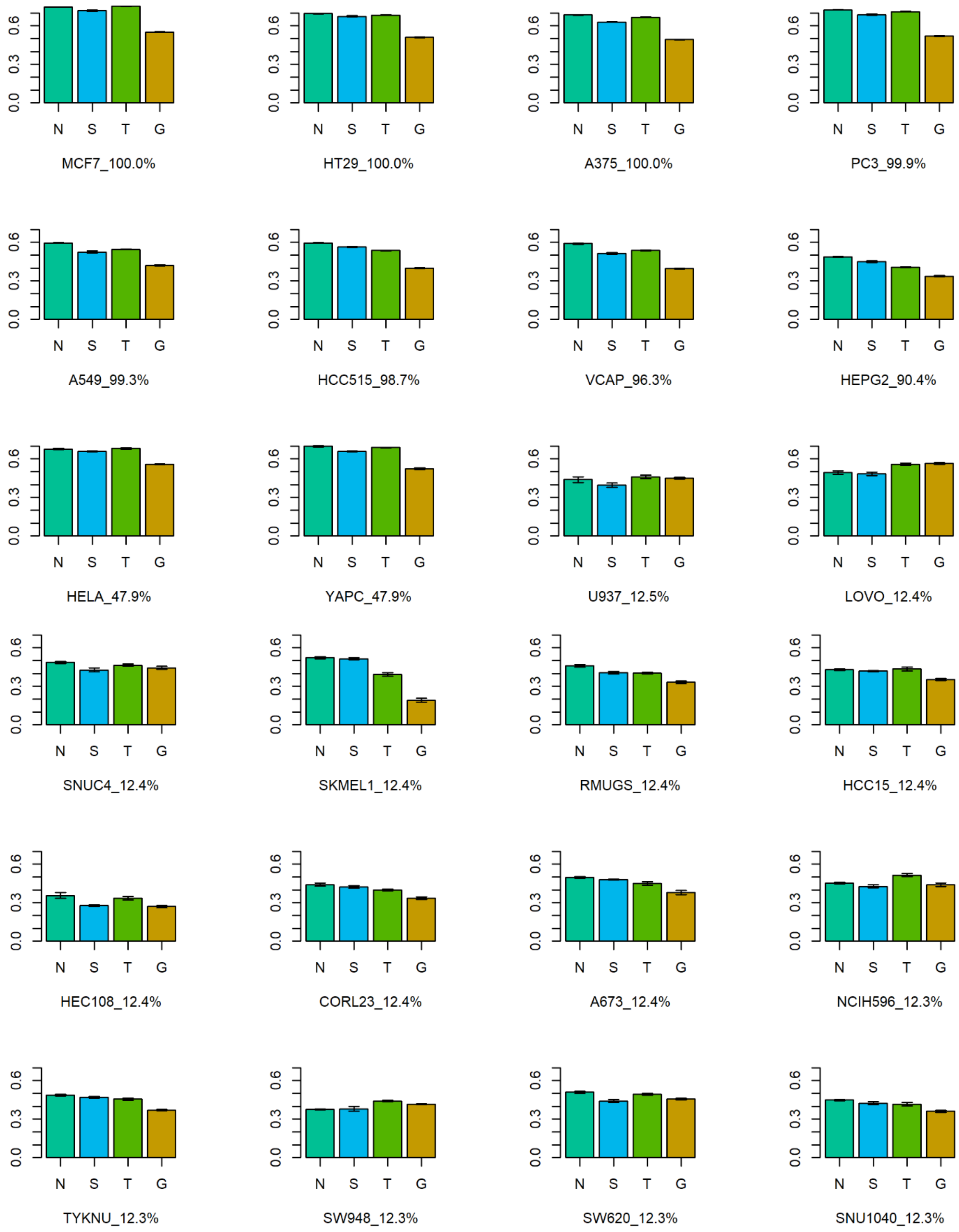
MCF7_100.0%
HT29_100.0%
A375_100.0%
PC3_99.9%
A549_99.3%
HCC515_98.7%
VCAP_96.3%
HEPG2_90.4%
HELA_47.9%
YAPC_47.9%
U937_12.5%
LOVO_12.4%
SNUC4_12.4%
SKMEL1_12.4%
RMUGS_12.4%
HCC15_12.4%
HEC108_12.4%
CORL23_12.4%
A673_12.4%
NCIH596_12.3%
TYKNU_12.3%
SW948_12.3%
SW620_12.3%
SNU1040_12.3%
N S T G
0.0 0.3 0.6

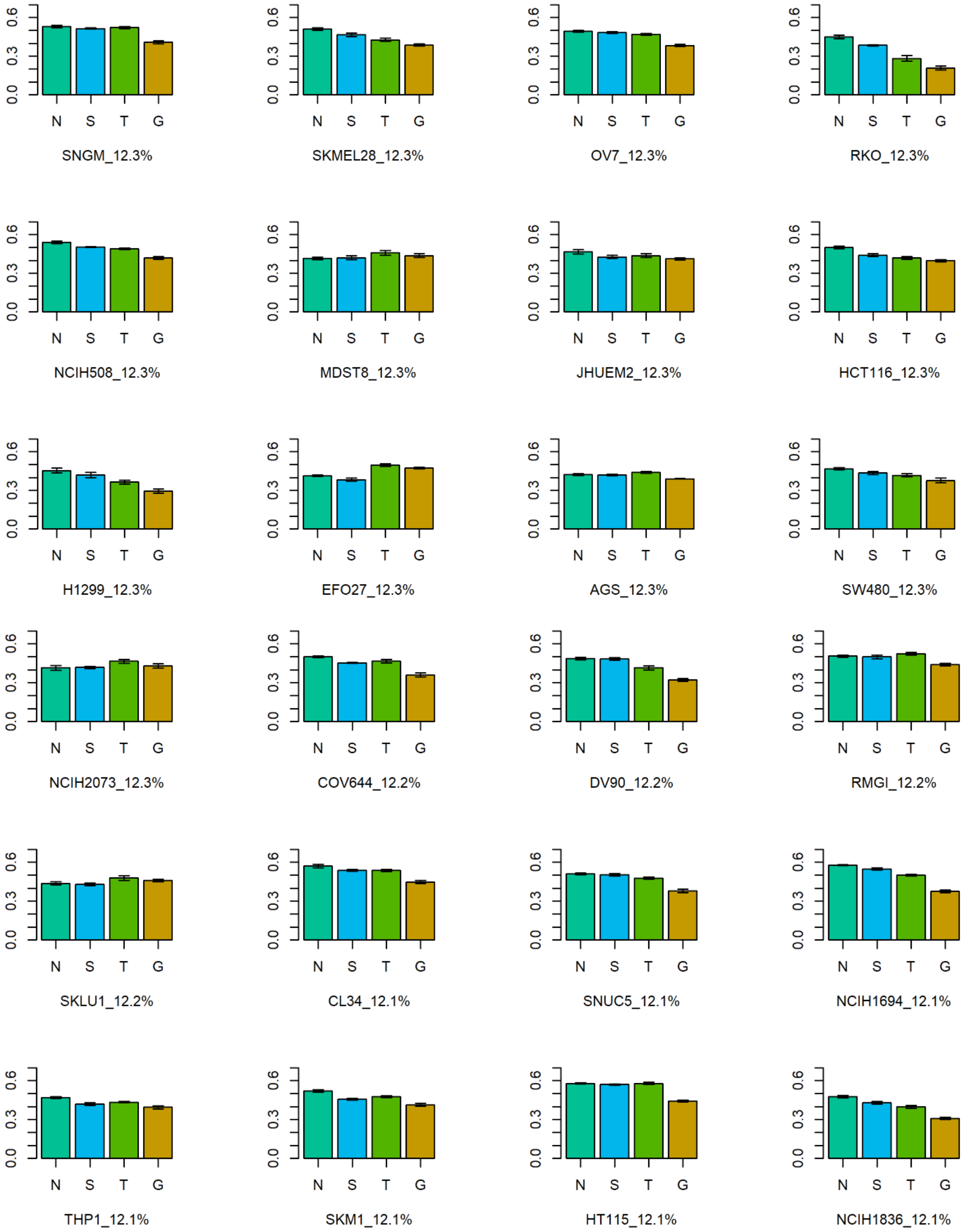
0.0
0.3
0.6
N
S
T
G
SNGM_12.3%
SKMEL28_12.3%
OV7_12.3%
RKO_12.3%
NCIH508_12.3%
MDST8_12.3%
JHUEM2_12.3%
HCT116_12.3%
H1299_12.3%
EFO27_12.3%
AGS_12.3%
SW480_12.3%
NCIH2073_12.3%
COV644_12.2%
DV90_12.2%
RMGI_12.2%
SKLU1_12.2%
CL34_12.1%
SNUC5_12.1%
NCIH1694_12.1%
THP1_12.1%
SKM1_12.1%
HT115_12.1%
NCIH1836_12.1%

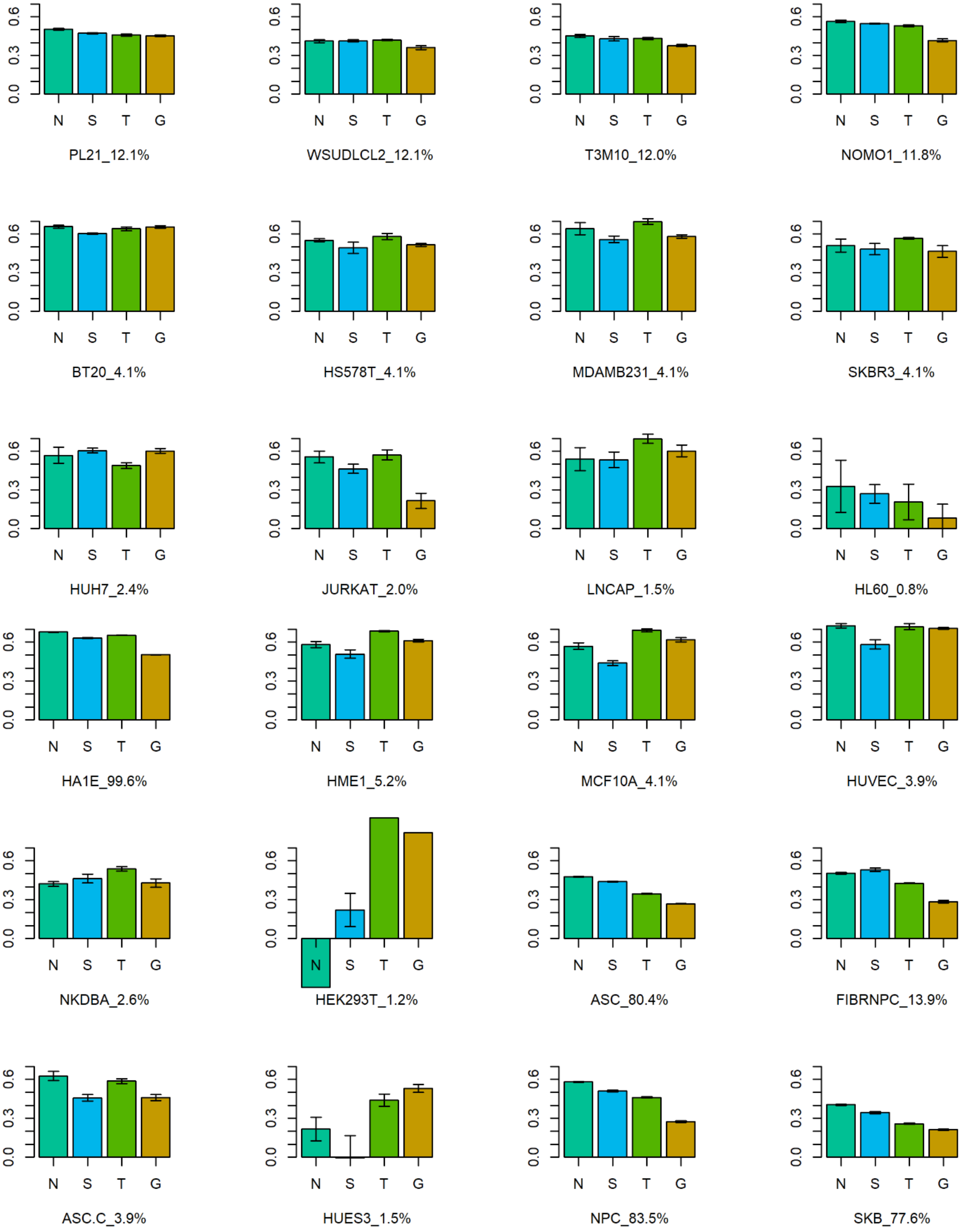
PL21_12.1%
WSUDLCL2_12.1%
T3M10_12.0%
NOMO1_11.8%
BT20_4.1%
HS578T_4.1%
MDAMB231_4.1%
SKBR3_4.1%
HUH7_2.4%
JURKAT_2.0%
LNCAP_1.5%
HL60_0.8%
HA1E_99.6%
HME1_5.2%
MCF10A_4.1%
HUVEC_3.9%
NKDBA_2.6%
HEK293T_1.2%
ASC_80.4%
FIBRNPC_13.9%
ASC.C_3.9%
HUES3_1.5%
NPC_83.5%
SKB_77.6%
N
S
T
G
0.0
0.3
0.6

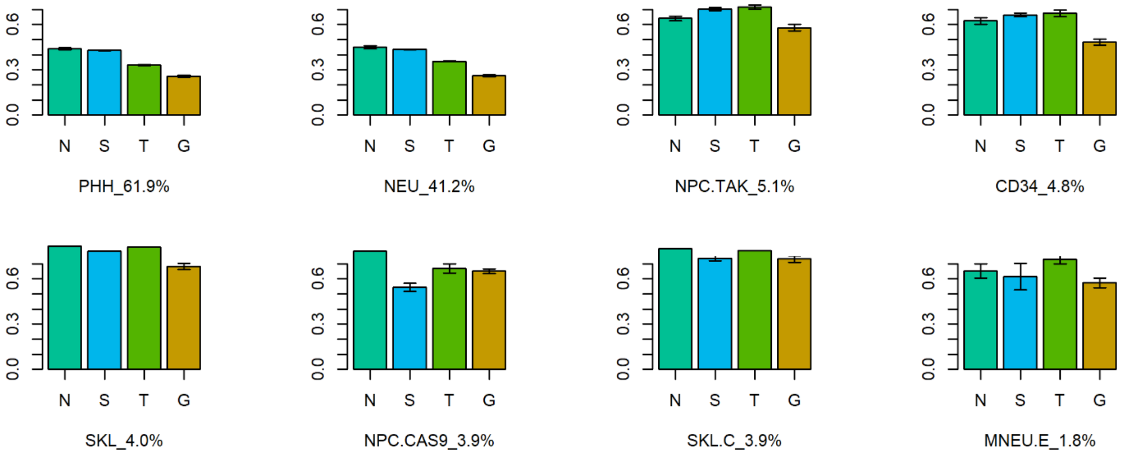


Figure 3: Negative weighted connectivity correlation across all genes and drugs, for each of the 80 cell lines in the sparse matrix. Methods are denoted by single-letter labels: N : neighborhood collaborative approach; S : SVD; T : two-way average; G : tissue-aGnostic (baseline method). Organized by cell type (cancer, immortalized, stem and primary) and ordered by percentage of drugs profiled in each cell. Error bars show variation across cross validation runs.

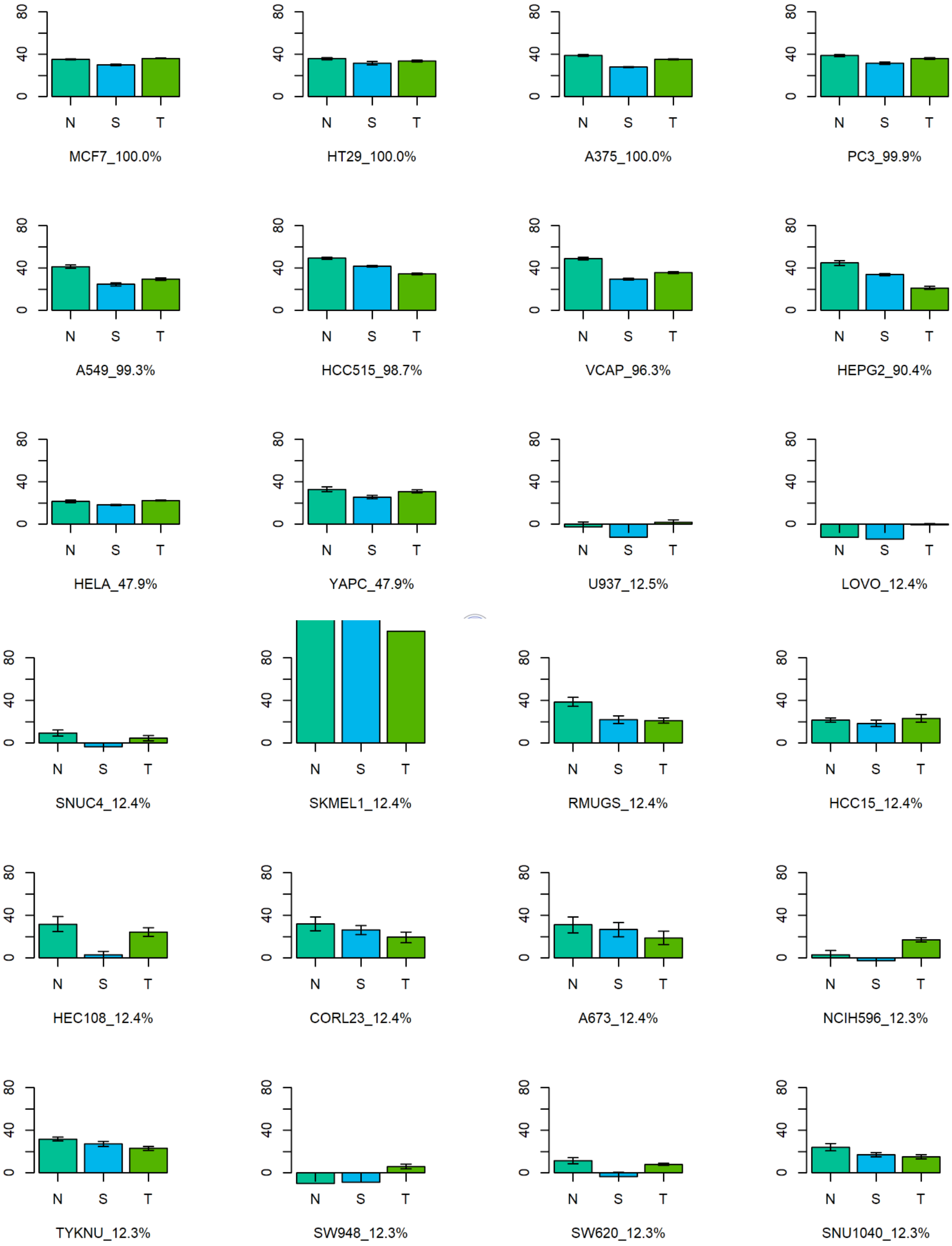
0
40
80
N
S
T
MCF7_100.0%
HT29_100.0%
A375_100.0%
PC3_99.9%
A549_99.3%
HCC515_98.7%
VCAP_96.3%
HEPG2_90.4%
HELA_47.9%
YAPC_47.9%
U937_12.5%
LOVO_12.4%
SNUC4_12.4%
SKMEL1_12.4%
RMUGS_12.4%
HCC15_12.4%
HEC108_12.4%
CORL23_12.4%
A673_12.4%
NCIH596_12.3%
TYKNU_12.3%
SW948_12.3%
SW620_12.3%
SNU1040_12.3%

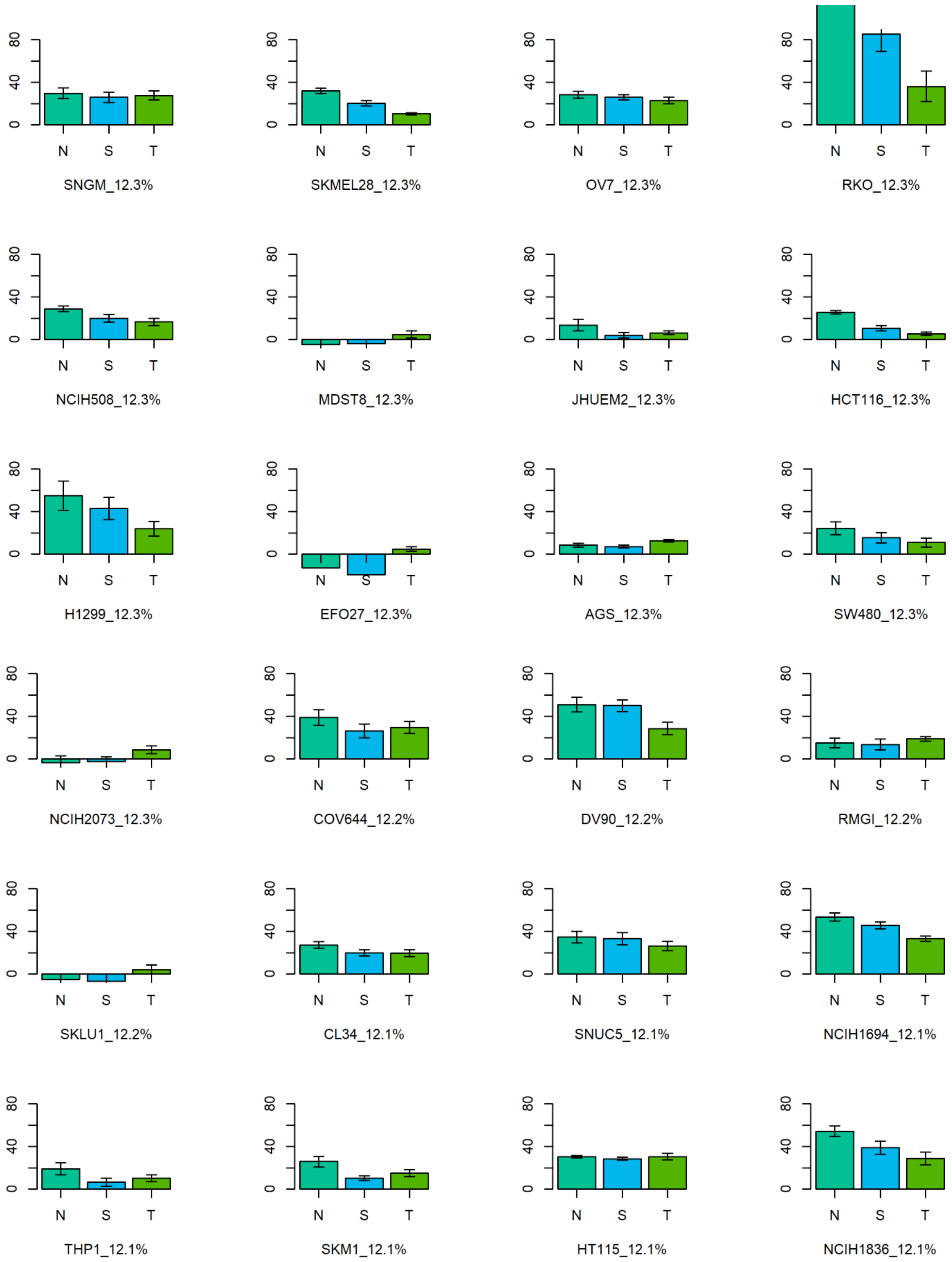
0
40
80
N
S
T
SNGM_12.3%
SKMEL28_12.3%
OV7_12.3%
RKO_12.3%
NCIH508_12.3%
MDST8_12.3%
JHUEM2_12.3%
HCT116_12.3%
H1299_12.3%
EFO27_12.3%
AGS_12.3%
SW480_12.3%
NCIH2073_12.3%
COV644_12.2%
DV90_12.2%
RMGI_12.2%
SKLU1_12.2%
CL34_12.1%
SNUC5_12.1%
NCIH1694_12.1%
THP1_12.1%
SKM1_12.1%
HT115_12.1%
NCIH1836_12.1%

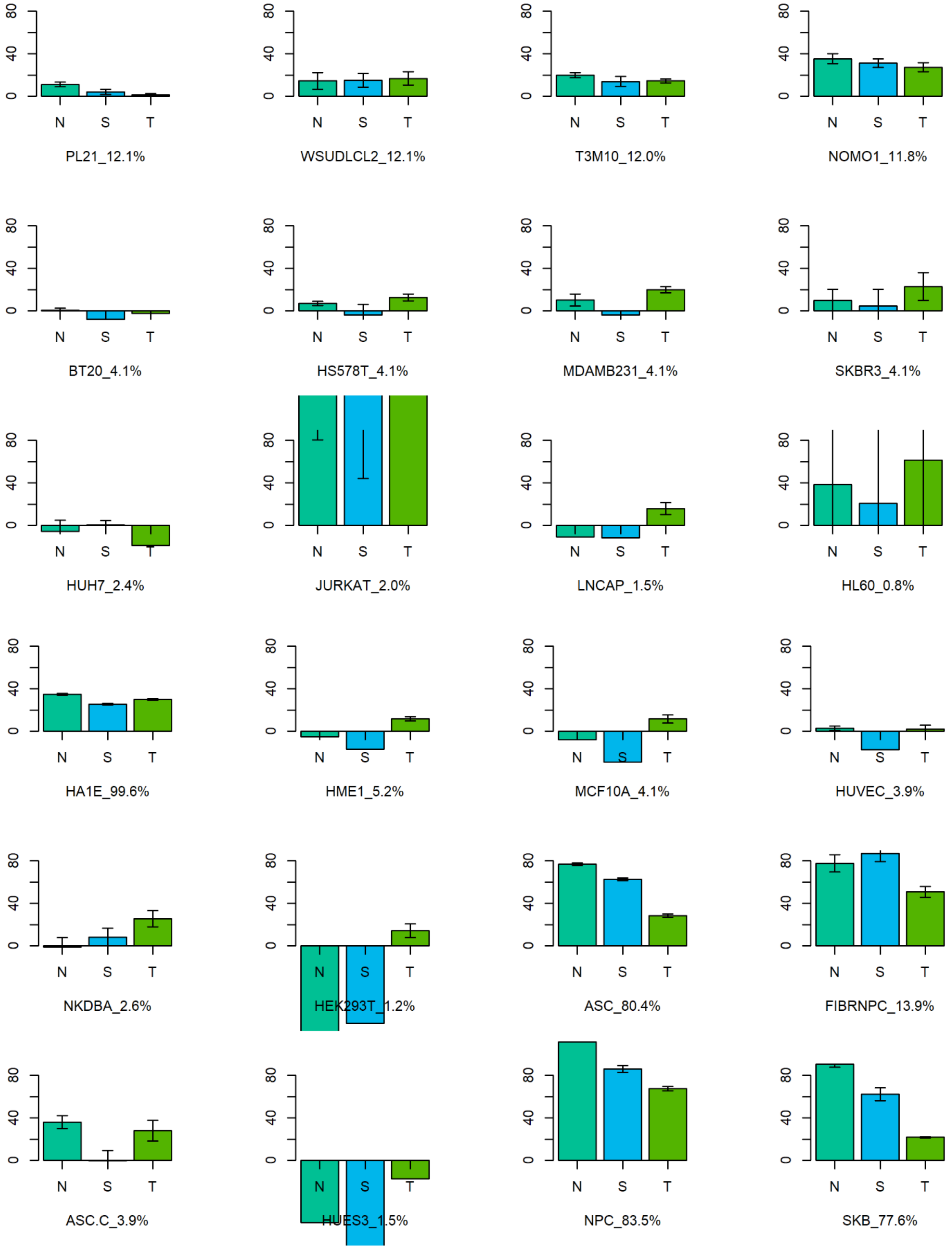
80
40
0
N
S
T
PL21_12.1%
WSUDLCL2_12.1%
T3M10_12.0%
NOMO1_11.8%
BT20_4.1%
HS578T_4.1%
MDAMB231_4.1%
SKBR3_4.1%
HUH7_2.4%
JURKAT_2.0%
LNCAP_1.5%
HL60_0.8%
HA1E_99.6%
HME1_5.2%
MCF10A_4.1%
HUVEC_3.9%
NKDBA_2.6%
HEK293T_1.2%
ASC_80.4%
FIBRNPC_13.9%
ASC.C_3.9%
HUES3_1.5%
NPC_83.5%
SKB_77.6%

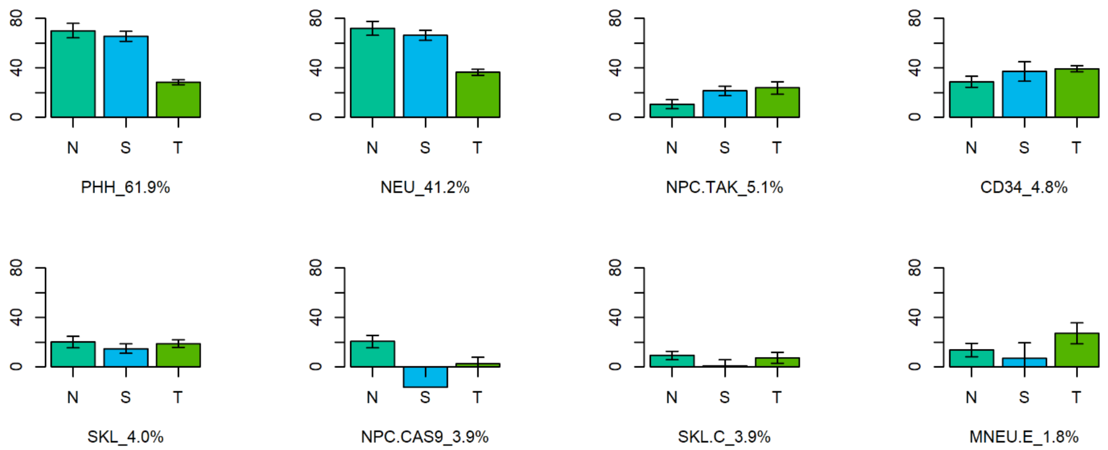


Figure 4: Percent change in negative weighted connectivity correlation compared to the tissue-agnostic method. Methods are denoted by single-letter labels: N : neighborhood collaborative approach; S : SVD; T : two-way average; G : tissue-aGnostic (baseline method). Organized by cell type (cancer, immortalized, stem and primary) and ordered by percentage of drugs profiled in each cell. Error bars show variation across cross validation runs.

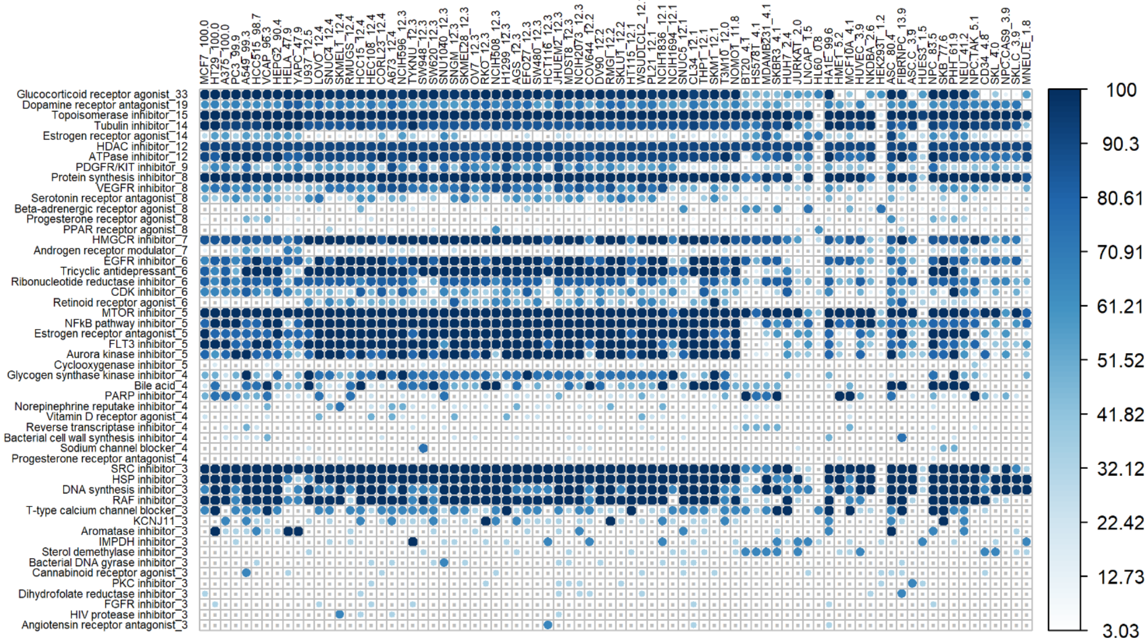

Figure 5: Percent of drugs correctly expressing strong connectivity to their drug class using the fully imputed sparse matrix by the neighborhood approach. Cells are organized by cell type (cancer, immortalized, stem and primary) and ordered by percentage of drugs profiled in each cell. PCL sets are ordered by the number of drugs in each set that are in the sparse matrix. The darker the shade of blue, the higher percentage of drugs with statistically significant NES scores. Grey dots represent PCL/cell combinations in which there were no statistically significant NES scores. In the main text, primary cells were pulled out and the plot was transposed for readability.

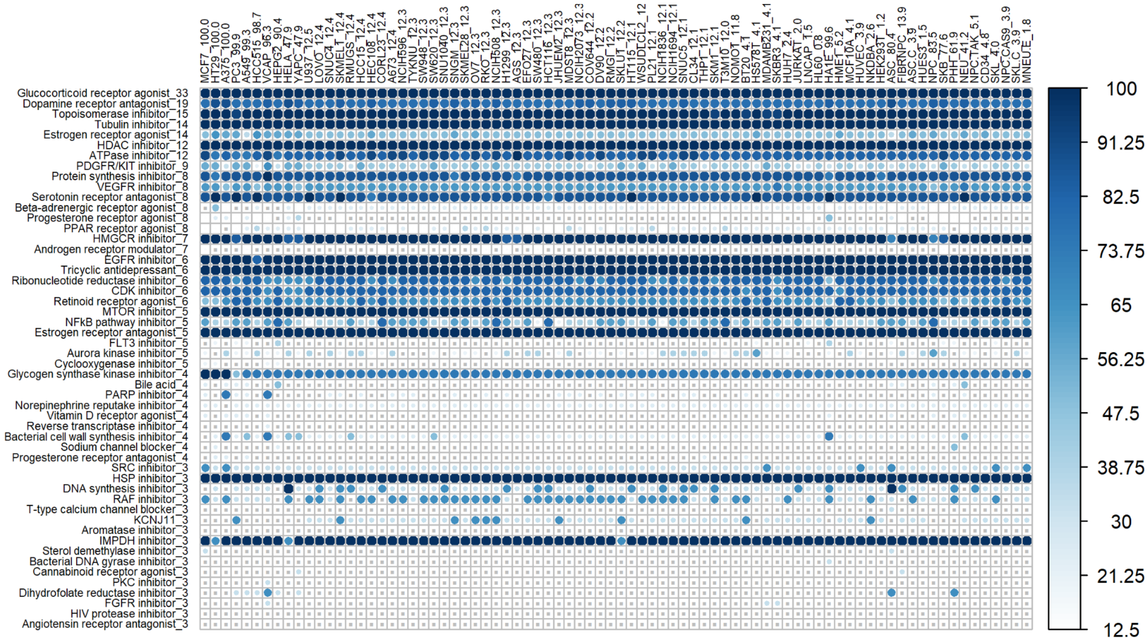


Figure 6: Percent of drugs correctly expressing strong connectivity to their drug class using the fully imputed sparse matrix by the tissue agnostic approach. Cells are organized by cell type (cancer, immortalized, stem and primary) and ordered by percentage of drugs profiled in each cell. PCL sets are ordered by the number of drugs in each set that are in the sparse matrix. The darker the shade of blue, the higher percentage of drugs with statistically significant NES scores. Grey dots represent PCL/cell combinations in which there were no statistically significant NES scores. In the main text, primary cells were pulled out and the plot was transposed for readability.

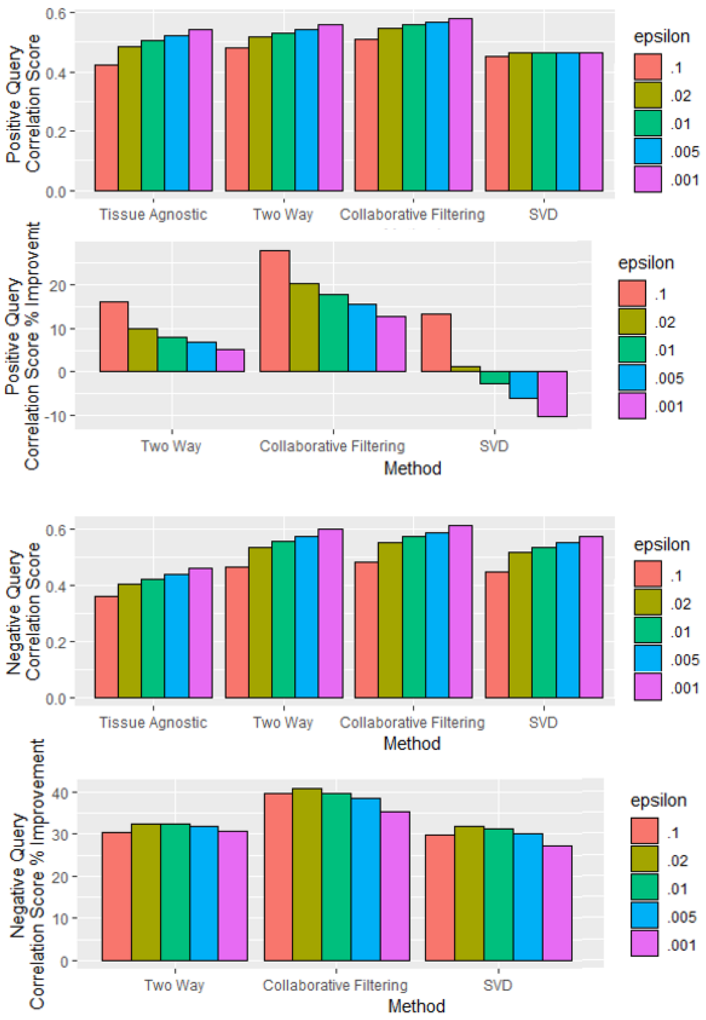


Figure 7: Change in average positive and negative query correlation scores across all cells and drugs obtained by varying $\epsilon$ by a factor of either two or ten in either direction from the values of $\epsilon = .01$ used in this work.

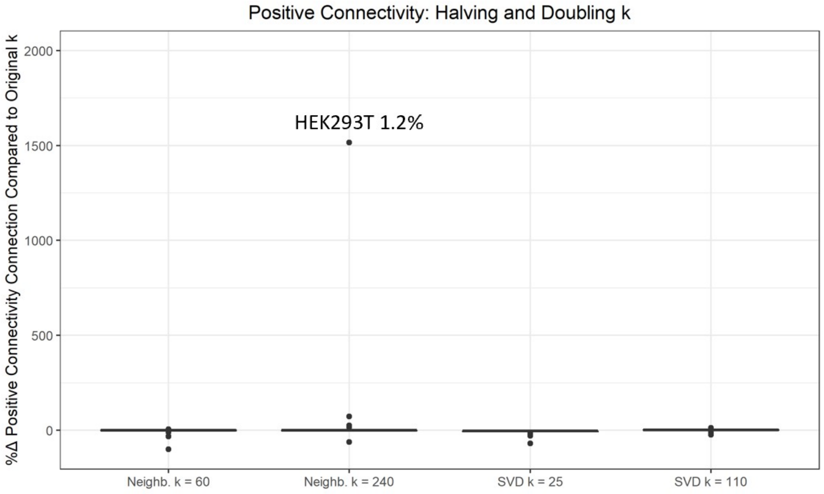


Figure 8: Percent change in average positive weighted connectivity correlation across all drugs and genes obtained by varying $k$ by a factor of two from the values of $k = 120$ for nearest neighbors and $k = 55$ for svd used in this work for the sparse data set. Outlier cell lines labelled with %data.

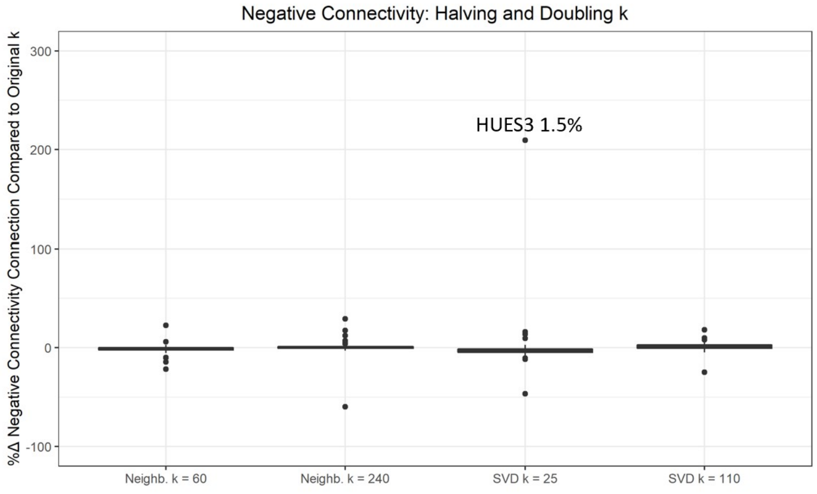


Figure 9: Percent change in average negative weighted connectivity correlation across all drugs and genes obtained by varying $k$ by a factor of two from the values of $k = 120$ for nearest neighbors and $k = 55$ for svd used in this work for the sparse data set. Outlier cell lines labelled with %data.
